# Depopulation of α-synuclein aggregates is associated with rescue of dopamine neuron dysfunction and death in a new Parkinson’s disease model

**DOI:** 10.1101/500629

**Authors:** Michal Wegrzynowicz, Dana Bar-On, Laura Calo’, Oleg Anichtchik, Mariangela Iovino, Jing Xia, Sergey Ryazanov, Andrei Leonov, Armin Giese, Jeffrey W. Dalley, Christian Griesinger, Uri Ashery, Maria Grazia Spillantini

## Abstract

Parkinson’s Disease (PD) is characterized by the presence of α-synuclein aggregates known as Lewy bodies and Lewy neurites, whose formation is linked to disease development. The causal relation between α-synuclein aggregates and PD is not well understood. We generated a new transgenic mouse line (MI2) expressing human, aggregation-prone truncated 1-120 α-synuclein under the control of the tyrosine hydroxylase promoter. MI2 mice exhibit progressive aggregation of α-synuclein in dopaminergic neurons of the substantia nigra pars compacta and their striatal terminals. This is associated with a progressive reduction of striatal dopamine release, reduced striatal innervation and significant nigral dopaminergic nerve cell death starting from 6 and 12 months of age, respectively. Overt impairment in motor behavior was found in MI2 mice at 20 months of age, when 50% of dopaminergic neurons are lost. These changes were associated with an increase in the number and density of 20-500nm α-synuclein species as shown by *d*STORM. Treatment with the oligomer modulator anle138b, from 9-12 months of age, restored striatal dopamine release and prevented dopaminergic cell death. These effects were associated with a reduction of the inner density of α-synuclein aggregates and an increase in dispersed small α-synuclein species as revealed by *d*STORM. The MI2 mouse model recapitulates the progressive dopaminergic deficit observed in PD, showing that early synaptic dysfunction precedes dopaminergic axonal loss and neuronal death that become associated with a motor deficit upon reaching a certain threshold. Our data also provide new mechanistic insight for the effect of anle138b’s function *in vivo* supporting that targeting α-synuclein aggregation is a promising therapeutic approach for PD.

## INTRODUCTION

Parkinson’s Disease (PD) and other α-synucleinopathies are characterized by aggregation of α-synuclein (αSyn) in Lewy bodies (LBs), Lewy neurites (LNs) and glial cytoplasmic inclusions (Jellinger and Lantos, 2010; Spillantini et al., 1998; Spillantini et al., 1997). It is now widely accepted that the process of αSyn aggregation in PD is directly involved in the pathogenesis and progression of the disease, and motor symptoms are mainly related to the dysfunction of the nigrostriatal dopaminergic (DA) system. Indeed, an impairment of DA neurotransmission in the striatum and loss of DA neurons in substantia nigra pars compacta (SNpc) are observed at the time of PD diagnosis (Hindle, 2010). To date, there is no treatment that affects the mechanism of the disease, and existing therapies are symptomatic.

Availability of an animal model of PD that reproduces progressive DA dysfunction and DA neuronal death with progressive αSyn aggregation in the nigrostriatal system is crucial for understanding disease mechanisms and the testing of novel therapies for treating movement deficit. In addition, until recently it was difficult to study quantitatively the early steps of αSyn aggregation and the effects of aggregate inhibitors in a living system, mainly due to the limitations in diffraction of conventional microscopy. Today, with the use of super resolution microscopy like STORM, dSTORM, STED and aptamer DNA PAINT (Sigal et al., 2018) and advanced analysis algorithm (Bar-On et al., 2012), it is possible to identify changes in αSyn aggregation at the single molecule level also in *in vivo* systems and understand the mechanistic nature of aggregate inhibitors.

Here we report a new transgenic mouse model (MI2 mice) expressing human 1-120 truncated αSyn that develops a progressive phenotype including αSyn aggregation, loss of striatal DA, reduction in induced striatal DA release, DA neuron death in SNpc, loss of nigro-striatal dopaminergic innervation, and motor deficits. We show that progressive DA dysfunction is associated with increased formation of synaptic striatal αSyn aggregates. Moreover, we show that the oligomer modulator, anle138b, previously shown to be effective in other models of protein aggregation including αSyn (Heras-Garvin et al., 2018; Martinez Hernandez et al., 2018; Wagner et al., 2015; Wagner et al., 2013), restores striatal DA release and prevents DA cell loss even when administration is started after the onset of DA dysfunction. For the first time and using *d*STORM and immunoblotting, we show that *in vivo*, in mouse brain, anle138b rescue of the DA deficit is associated with reduction of the density of large αSyn aggregates and an increase in dispersed monomeric and small assemblies of 1-120hαSyn.

## MATERIALS AND METHODS

### Study design

This study aimed to generate and characterize a novel transgenic mouse model of PD that would enhance our understanding of the relationship between progressive αSyn aggregation and mechanisms of disease and allow testing prospective therapies for α-synucleinopathies. The homozygous line expressing 1-120hαSyn under the TH promoter, in a null endogenous αSyn background was generated and used for the experiments. The αSyn-null C57Bl/6S strain was used as a control line. C57Bl/6J mice, expressing endogenous mouse αSyn were also included in the experiments where transgenic protein expression was compared to physiological mouse αSyn. Moreover, DA characteristics in MI2 mice were compared with both C57Bl/6S and C57Bl/6J mice to confirm that differences between C57Bl/6S and MI2 mouse lines were due to expression of the transgene rather than absence of αSyn in the C57Bl/6S animals.

MI2 and control mice were obtained from separate lines, therefore their genotypes were known at the time of experimental design. However, for behavioral experiments and anle138b-treatment mice were assigned randomly to the experimental groups. For each experiment, mouse numbers and statistical tests are described in the figure legends and in Supplemental Information. Analysis of dopaminergic impairment, namely measurements of DA in striatal homogenates and microdialysates, stereological counting of nigral cells and striatal innervation were performed by researchers blinded to experimental conditions.

### Ethics

This research has been performed under the Animals (Scientific Procedures) Act 1986 Amendment Regulations 2012 following ethical review by the University of Cambridge Animal Welfare and Ethical Review Body (AWERB), under project license no. 70/8383.

### Generation of transgenic mice

The MI2 mouse line was generated using the same procedures as described by Tofaris et al. for α-Syn120 mice (Tofaris et al., 2006). Briefly, a transgene construct containing human truncated 1-120 αSyn, subcloned downstream of the rat TH promoter (Figure 1A) was injected into αSyn-null C57Bl/6OlaHsd (C57Bl/6S) mouse (Specht and Schoepfer, 2001) pronuclei. The presence of the transgene was detected in the founders and progeny by PCR. Founders were then bred with αSyn-null C57Bl/6S mice and mouse αSyn-negative/1-120hαSyn-positive progeny were crossed to homozygosity.

**Figure 1.**
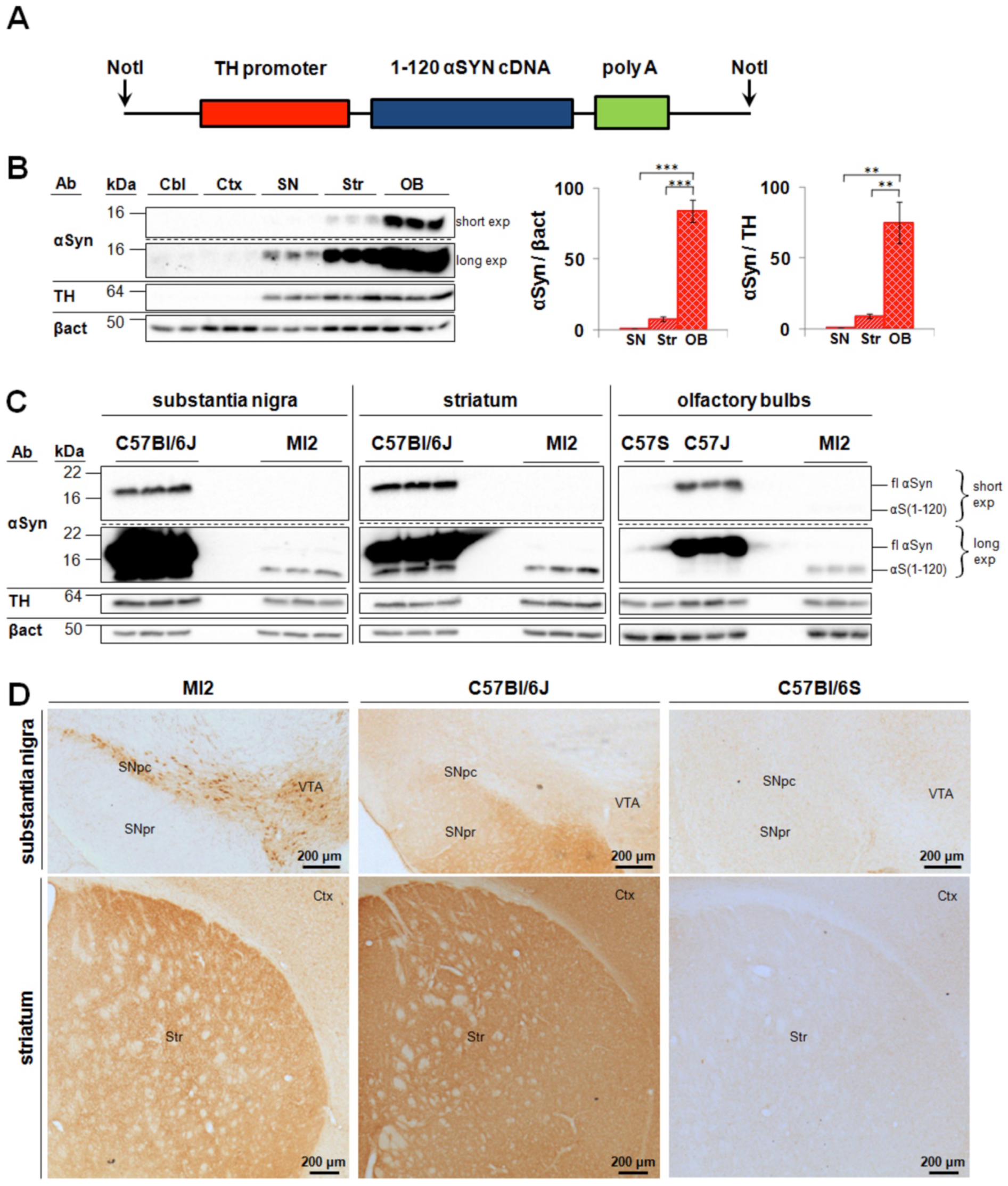
Expression of transgenic 1-120 hαSyn in MI2 mice. (A) 1-120 hαSyn transgene construct used for generating the MI2 mice. (B) Immunoblots showing αSyn in lysates from cerebellum (Cbl), cortex (Ctx), substantia nigra (SN), striatum (Str) and olfactory bulbs (OB) of 1.5 month-old mice following short (short exp) and long (long exp) exposure times. Quantification was performed for SN, Str and OB, but not for Cbl and Ctx, where the signal was negligible. Data are expressed as fold difference compared to SN (mean ± SEM, n=3 mice, one-way ANOVA with Bonferroni correction (**p<0.01, ***p<0.001) (Detailed statistics in Suppl. Material). (C) Immunoblots comparing expression levels between control and MI2 mouse lines. Expression of 1-120 hαSyn in MI2 mice is much lower than that of endogenous full-length αSyn (fl) in WT C57Bl/6J in the OB. Interestingly some truncated αSyn with a similar size to the transgenic 1-120 αSyn can be seen in protein extracts of SN and Str of WT C57Bl/6J mice. (D) Immunohistochemistry of brain sections of 1.5 month-old MI2 mice detected with the Syn1 antibody shows 1-120hαSyn protein in neuronal cell bodies and processes in SNpc and ventral tegmental area (VTA) and in neuropil in striatum (see also Figure S1A). In C57Bl/6J mice, endogenous αSyn, is also found in SN pars reticulata (SNpr) and Ctx besides SNpc, VTA and Str. The specificity of Syn1 antibody for αSyn was confirmed by the absence of staining in C57Bl/6S mice that lack endogenous αSyn.

### Western blotting

Animals were sacrificed by cervical dislocation, brains were snap frozen on dry ice and kept at -80°C. Brain regions were isolated, homogenized in ice-cold PBST (1 × PBS (Gibco), 0.3% Triton X-100 (Sigma), protease inhibitor cocktail (Roche)) and centrifuged at 4°C, at 14 000 × g, for 15 min. Supernatant protein content was measured using a BCA protein assay kit (Novagen) and protein concentration normalized using PBST. Following 5 min at 95°C in 3 × Laemmli buffer, samples were separated by SDS-PAGE, blotted onto nitrocellulose membranes (Bio-Rad) and proteins were crosslinked to the membrane with 4% paraformaldehyde (PFA) for 30 min. Non-specific background was blocked with 5% milk in TBST and incubated overnight at 4°C with primary antibodies (mouse anti-αSyn (Syn1), BD Biosciences, 1:500; rabbit anti-TH, Abcam, 1:1000; rabbit anti-β-actin, Abcam, 1:10000) in 5% milk. Membranes were then incubated with peroxidase-conjugated secondary antibodies (GE Healthcare or DAKO, 1:5000) and the blots imaged using a Chemi Doc MP imager (Bio-Rad), using West Dura Extended Duration Chemiluminescent Substrate (Thermo Fisher Scientific). Blots were analyzed using Image Lab 5.1 (Bio-Rad Laboratories).

Bands corresponding to mouse full length monomeric (∼17 kDa) and 1-120 truncated human αSyn (∼14 kDa) were observed in C57Bl/6J and MI2 mice respectively. No αSyn was present in C57Bl/6S (Figure 1C). Non-specific bands recognized by the Syn1 antibody were previously reported (Figure 2C) (Perrin et al., 2003).

**Figure 2.**
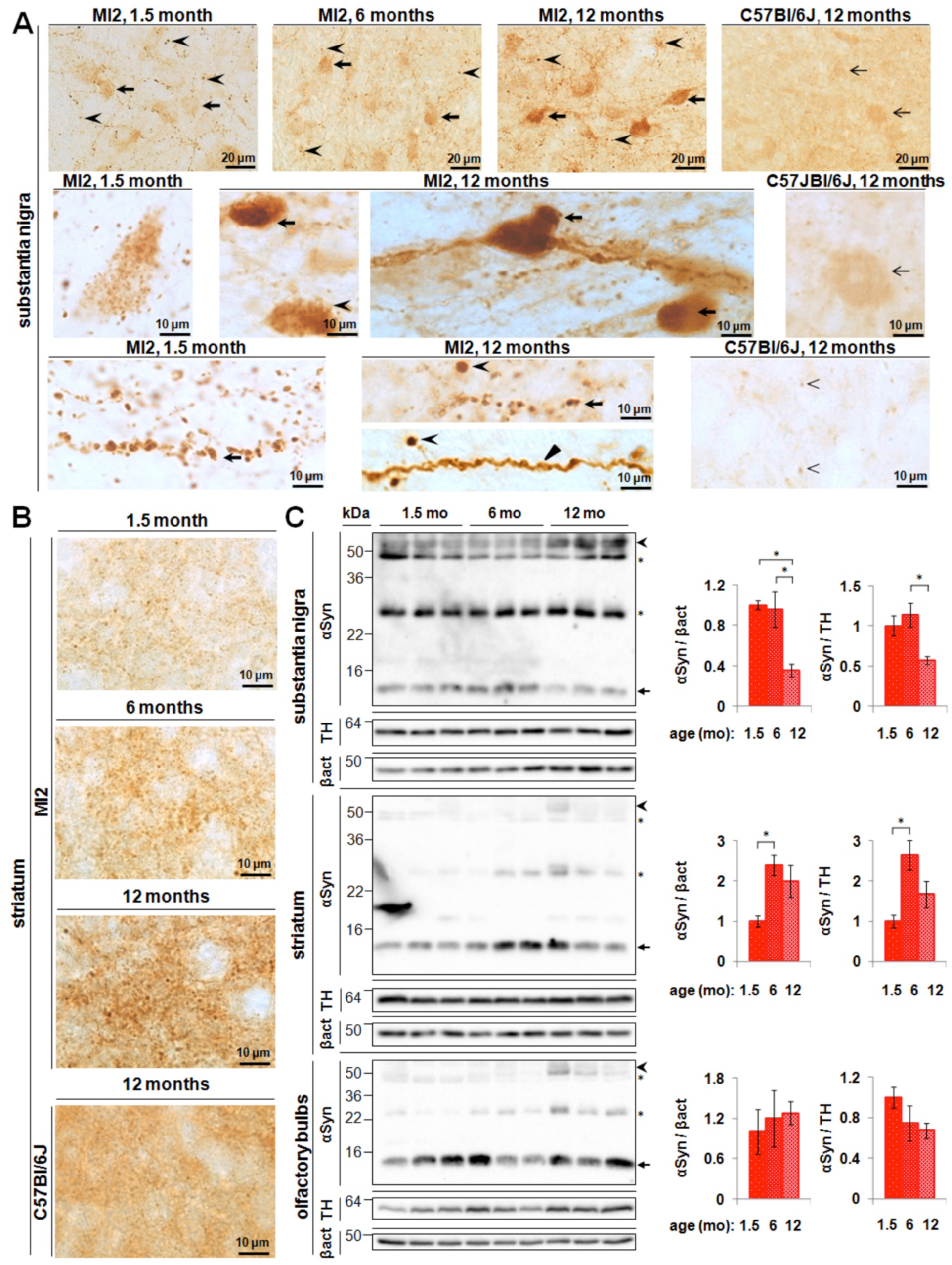
Aggregation of 1-120hαSyn protein in MI2 mice. (A) Top panel: progressive accumulation of 1-120 hαSyn protein with age in SNpc of MI2 mice in cell bodies (arrows) and processes (arrowheads). Middle panel: abundant small inclusions of 1-120hαSyn protein in SNpc cell bodies are present at 1.5 months and large, LB-like aggregates in SNpc neurons (arrows) among cells with 1-120hαSyn punctate staining (arrowhead) are found at 12 months of age. Bottom panel: large 1-120hαSyn puncta are distributed along the processes in SNpc at 1.5 and 12 months of age (arrows). At 12 months of age, processes uniformly filled with 1-120 hαSyn (triangle) and round inclusions containing condensed 1-120 hαSyn protein (arrowheads) are also visible. The staining of αSyn in WT C57Bl/6J mice is much less intense and more homogenous. Middle right panel: less αSyn is present in cell bodies, and no cellular inclusions are found (arrow). Bottom right panel: αSyn puncta in the nigral processes are much less numerous and smaller than in MI2 mice (arrowheads). (B) Progressive accumulation of 1-120 hαSyn puncta in MI2 striatal neuropil in 1.5, 6 and 12 month-old MI2 mice. In 12 month-old C57Bl/6J mice striatal αSyn is distributed more homogenously and large αSyn-positive puncta are not present (bottom panel). (C) Immunoblotting of brain lysates from MI2 mice as a function of age. The levels of monomeric 1-120 hαSyn (∼14 kDa, arrow) shown in the western blots were quantified and normalized to either β-actin or TH (right panels). Data are expressed as fold difference compared to 1.5 month-old animals (mean ± SEM, n=3 mice, one-way ANOVA, multiple comparison with Bonferroni corrections). In SN, a reduction of monomeric 1-120hαSyn was found as a main effect of age (statistically significant differences between 1.5 and 12 months and between 6 and 12 months (αSyn/β-actin or αSyn/TH (*p<0.05) for all comparisons. In the striatum there was a significant increase of 1-120hαSyn between 1.5 and 6 months of age (*p<0.05) for both αSyn/β-actin and αSyn/TH. There was no significant change in OB. In all the three regions increased amounts of high molecular weight (HMW) 1-120hαSyn bands (∼55 Da, arrowhead) were present in MI2 mice at 12 months compared to 1.5 and 6 months of age. Stars denote the non-specific bands recognized by Syn1 antibody in both MI2 and αSyn-null C57Bl/6S mice (Perrin et al., 2003). (See detailed statistical evaluation in Suppl. Material).

### Immunohistochemistry

Mice were anesthetized by an intraperitoneal administration of pentobarbital (Merial Animal Health), and transcardially perfused with ice-cold PBS, followed by 4% PFA in PBS. Brains were post-fixed overnight in 4% PFA, then transferred to 30% sucrose in PBS with 0.1% NaN_3_ (Sigma) and stored at 4°C. Brains were frozen and 30 µm sections cut using a freezing microtome (Bright). Endogenous peroxidase activity was inhibited by treatment with 3% H_2_O_2_ containing 20% methanol in PBST. Sections were blocked with 5% normal horse serum (NHS, Vector) in PBST. Antigen retrieval was performed before Syn1 staining, using 10 mM sodium citrate buffer pH 8.5 at 80°C for 30 min. Free floating sections were incubated overnight at RT with primary antibodies diluted in PBST (Syn1, 1:500; rabbit or chicken anti-TH, Abcam, 1:1000; mouse anti-NeuN, Millipore, 1:1000, rabbit anti-VAMP2, Abcam, 1:250) and then with biotinylated (Vector, 1:2000) or Alexa Fluor-conjugated (Thermo Fisher Scientific, 1:500) secondary antibodies diluted in PBST containing 5% NHS. Peroxidase-based staining was developed using Vectastain Elite ABC HRP Kit (Vector) and DAB Peroxidase Substrate Kit (Vector). Sections were mounted on microscope slides dehydrated, cleared in xylene and coverslipped using DPX Mountant (Sigma). (Thermo Fisher Scientific). In some experiments, sections were counterstained with 0.1% cresyl violet (Sigma). For immunofluorescence nuclei were counterstained with DAPI (0.5 µg/ml; Roche) and sections coverslipped using FluorSave mounting medium (Calbiochem). DAB-stained sections were analyzed using Olympus BX 53 light microscope and fluorescent sections using a Leica DMI 4000B epifluorescent microscope.

For dSTORM, 30 µm free floating striatal slices were stained using a similar protocol, but PBST containing 0.25% Tween 20 was used and the non specific background blocking solution contained 5% goat serum and 1% bovine serum albumin. Sections were stained with Syn204 mouse anti-human αSyn (abcam, 1:250) and Alexa Fluor 647 secondary anti-mouse antibody (Thermo Fisher Scientific, 1:500). In some experiments striatal sections were co-stained with rabbit anti-synaptobrevin 2 (VAMP2) antibody (Synaptic Systems, 1:500), followed by Cy3B anti-rabbit secondary antibody (1:250) to confirm synaptic localization of 1-120hαSyn by dSTORM.

### dSTORM

dSTORM images were taken using the Vutara microscope as described in (Bar-On et al., 2012), at power of 40% 647 and 2% 405 laser, and 3000 frames were taken for each movie. Localizations were gathered following the movie with a ratio of 10 of background to noise. All movies were then post analyzed using a denoise level of 0.5.

Cluster analysis, mean diameter cluster analysis and the number of localizations per clusters were determined using the density based algorithm (dbscan) followed by 2D principal component analysis and outliers detection as described in (Bar-On et al., 2012) (See Figure S2 for more details). For anle138b vs placebo treatment analysis, clusters in both conditions were additionally filtered according to inner density threshold determined according to the placebo condition per imaging day to avoid large-homogenous distributions that could form artifact clustering. Mean density of cluster was calculated for the placebo condition on individual day of measurement and clusters with less than 10% of this density were excluded from analysis in both placebo and anle138b treatment conditions measured on the same day. The localizations that were accounted for very low density clusters were in both cases added to the free (unclustered) populations of the proteins.

#### Monomeric αSyn definition and characterization

In order to characterize the dimensions of monomeric α-Syn by dSTORM, we plated/stripped a solution containing full-length recombinant human α-Syn on the same coverslips used for the brain slices and stained the solution following the same protocol and primary and secondary antibodies, used for the striatum slices. We imaged the recombinant proteins using dSTORM, and analyzed the size (mean diameter size, Figure S5A) and density of each one of the recombinant proteins particles using density-based (dbscan) cluster analysis (Figure S2 and Figure S5). We first used imageJ to identify putative centers of mass for each recombinant protein by loading all dSTORM images of the proteins (Figure S5B). We removed very high or very low frequencies, to easily locate local intensity maxima, then we performed a reverse Fourier transform and used a build-in function in the program of tracking local maxima using a threshold of signal to noise to assure the maxima and avoid artifacts (Figure S5B). The tracked local maxima were then used as a reference to compare the tracking of putative clusters by the dbscan (density based algorithms) as was used for the slices analysis. Several parameters of ε (size of area for search) and k (threshold number of neighbors) were tested and compared to the results obtained from the imageJ. Finally the most accurate set of parameters for the dbscan was found to be ε=50 nm and k=4 neighbors (Figure S5B). Based on analysis of 870 putative recombinant proteins in 15 different images, the final analysis indicated that the size of a single monomeric recombinant protein could correspond to a median size of 20.6 nm including a median number of 7 fluorophores per protein (Figure S5A).

#### *Analysis of the non-clustered* αSyn *population following anle138b treatment*

Using the definition and parameters of the monomeric recombinant αSyn analysis as described above, we characterized the population of αSyn proteins that were not defined as aggregates (non-clustered) in the mice treated with anle138b. The goal was to try to explore what the characteristics of this population are, whether the population is comprised mainly of monomers-like or higher orders of protein clusters/aggregates. When comparing anle138b-treated mice with placebo-treated animals, we found that the non-clustered population of proteins was substantially higher, 53.4% for anle138b-treated mice vs 23% for placebo-treated animals (Figure 7D). We also noticed that among the population of non-clustered proteins there were aggregates with very low density, where the proteins in the aggregate seem dispersed (their density was defined as at least 10 times lower than the average density of aggregates from placebo-treated MI2 mice). In order to characterize more deeply these population of dispersed aggregates, we analyzed only this population of non-aggregated protein using the same parameters found for estimating recombinant monomeric αSyn as indicated above and we found that over 80% of the population is defined in a very similar way as a monomeric protein with a median diameter of 23nm and median of 7 fluorophores per single cluster (protein). When looking at the overall distribution we can observe that the dispersed assemblies are composed of smaller assemblies with a mean diameter of 50-100 nm, that corresponds only to 10% of the total protein clusters (Figure S5C) which were surrounded or bridged to each other by what we had estimated as monomeric αSyn (around 85% of the total clusters, Figure S5C). In Figure S5D, a representative example of one of these dispersed assemblies is shown – with cores of smaller aggregates that are denser with a mean diameter size>50 nm and surrounded by many monomer-like species (mean diameter size=23 nm).

### Dopamine measurement

Bilateral striata were isolated from frozen brains, weighed and homogenized in 0.2 M perchloric acid. Homogenates were centrifuged at 6000 × g for 20 min at 4°C. The supernatants were resolved by high performance liquid chromatography with a Hypersil BDS C18 reversed phase column (3 µm particle size, 130 Å pore size, 100 × 4.6 mm; Phenomenex) at a flow rate of 1 ml/min as previously reported (Garcia-Reitbock et al., 2010). The mobile phase comprised citric acid (31.9 g/l), sodium acetate (2 g/l), octanesulfonic acid (460 mg/l), EDTA (30 mg/l) and methanol (15%), pH 3.6. DA was detected by redox oxidation using an ESA 5014 analytical cell and ESA Coulochem II electrochemical detector (Thermo Fisher Scientific) with reducing (E1) and oxidizing (E2) electrodes set at -200 mV and +250 mV, respectively. The chromatograms were analyzed using the Chromeleon Chromatography Data System (V 6.2; Dionex). DA levels were normalized to the initial wet tissue weight and expressed as pmol/mg.

### *In vivo* microdialysis

Animals were anesthetized with Caprieve and isoflurane, and placed in a stereotaxic frame under general anaesthesia. A craniotomy was drilled in the skull according to the following coordinates relative to bregma: +0.8 anteroposterior, +2.1 lateral (Paxinos and Franklin, 2004). The guide cannula CMA7 (CMA Microdialysis) was implanted in the brain, so that its tip reached the calloso-striatal border, -2.3 mm dorsoventral relative to the skull surface (Paxinos and Franklin, 2004), and was secured to the skull using glass ionomer luting cement FujiCEM 2 (GC Corporation) and two anchor screws CMA7431021 (CMA Microdialysis). Microdialysis was performed 24h following the surgery, when the mice had recovered. A microdialysis probe CMA7 (0.24 mm × 2 mm membrane, 6 kDa cut-off; CMA Microdialysis) was inserted through the guide cannula, and freely-moving mice were injected with artificial cerebro-spinal fluid (ACSF; 140 mM NaCl, 7.2 mM glucose, 3 mM KCl, 1 mM MgCl_2_, 1.2 mM CaCl_2_, 1.2 mM Na_2_HPO_4_, 0.27 mM Na_2_HPO_4_, pH 7.4) at 2 µl/min. An equilibration period of 30 min was allowed to recover basal levels of extracellular DA in striatum, and then dialysates were collected every 20 min in ice-cold tubes containing 5 µl of 0.2 M perchloric acid. After 60 min, ACSF was replaced with high-potassium ACSF (93 mM NaCl and 50 mM KCl), to induce neurotransmitter release, and a further 3 samples were collected, after which the physiological ACSF was infused again. Each sample was snap frozen in dry ice and stored at -80°C until HPLC analysis was performed as described above. Mice were sacrificed and the microdialysis probe position was verified in the striatum. For each individual mouse, DA release was normalized to the baseline fraction (0 min) and expressed as fold difference relative to the average DA release directly following K^+^ stimulation (60 min fraction) in the respective control C57Bl/6S group. DA release was compared between MI2 and C57Bl/6S mice and between anle138b- and placebo-treated mice.

### Stereological counting of nigral cells

Six 30 µm-thick coronal sections, evenly spaced at 180 µm within the range covering the whole SNpc (between -2.8 mm and -3.85 mm AP from bregma (Paxinos and Franklin, 2004)), were investigated in each mouse. TH immunohistochemistry was performed as described above, and tissue sections were analyzed with the optical fractionator probe using Stereo Investigator 11.07 (MBF Bioscience) with Olympus BX53 microscope equipped with QImaging Retiga camera and X-Y step motorized stage controlled by MAC 6000 System controller. The SNpc area was outlined under a 4xobjective and cells were counted under a 100xobjective using the following parameters: counting grid size - 220 × 220 µm, counting frame - 85 × 85 µm. The thickness of the section was measured at each counting site. Data were processed using built-in software and the optical fractionator formula to estimate the total number of TH-positive neurons in SNpc.

In order to confirm neurodegeneration in SNpc of MI2 mice, stereological counting of all nigral neurons was performed after staining the sections with the NeuN pan-neuronal marker. In order to contour SNpc for NeuN cell counting, TH immunofluorescence was performed in the same sections and staining visualized with Alexa Fluor 488 secondary antibody. Sections were visualised at 488 nm using an epifluorescent microscope and the SNpc was outlined based on TH immunofluorescence. The resulting contour was super-imposed in the bright field image of NeuN immunohistochemistry. The slides were transferred to a stereology microscope, and the contour of SNpc was manually reconstructed in bright field in Stereo Investigator 11.07 before stereological counting of NeuN-positive cells (Figure S4).

### Measurements of TH^+^ striatal innervation

Density of dopaminergic fibers in the striatum was estimated by the modified spherical probe counting method (Mouton et al., 2002) in two 30 µm-thick coronal sections, representing medial and caudal striatum (+0.12 mm and -0.66 mm AP from bregma (Paxinos and Franklin, 2004)). The tissue was processed for TH immunohistochemistry as described above, and analyzed using Stereo Investigator 11.07. A series of hemispherical (10 µm radius) probes, arranged in a 2 dimensional array, spaced at 250 µm intervals was superimposed on the striatum, and the total number of TH-positive fibers that cross the surface of the probe was counted under a 100x objective. The thickness of the section was measured at each counting site. The estimated total length of TH-positive fibres in individual sections was normalized to the measured volume of the section and averaged between counting sites. Data are expressed as nm of length of TH-positive fibers in µm^3^ of striatal volume.

### Rotarod test

Motor performance was tested using an accelerating rotarod (Ugo Basile). The test was performed in mice at 6, 12, 15 and 20 months of age using separate cohorts of animals at each time point. Both males and females were tested in MI2 and C57Bl/6S groups, but we observed that some 15 and 20 month-old MI2 females were overweight, therefore, since the body weight may affect motor performance (Kudwa et al., 2013; McFadyen et al., 2003), the heaviest individuals from this group were excluded from the test, to obtain no significant difference between the MI2 and C57Bl/6S females mice tested (t-test, p=0.552 at 15 months, p=0.851 at 20 months). Prior to the test, mice were trained for two days. Each training session consisted of four trials at constant speed of 16 rpm, for a maximum of 60s per trial. Mice were then tested using the acceleration mode (5-40 rpm) for three consecutive days with three trials per day and with at least 30 min intervals between the trials. Mice were placed on the rod and the time that the individual mouse took to fall from the rod was measured. Single trials were considered as technical replicates, and an average of 9 technical replicates (3 days per 3 trials) was calculated for each animal to obtain each biological replicate that was then used for statistical purposes.

### Static rod test

The cohort of 20 month-old mice used for the rotarod test were then used for the static rod test performed according to the method of Deacon with modifications (Deacon, 2013). Two 60 cm-long rods of 25 mm and 15 mm diameters were attached to the platform and held horizontally 30 cm above the cushioned surface. Mice were placed at the end of the rod, facing away the platform and were allowed to freely turn back and walk to the platform. Prior to the test animals received training trials using 25 mm and 15 mm rods. On the next day mice were tested on both 25 mm and 15 mm rods (one trial per each diameter). Two parameters were measured: orientation time (time taken to turn from the initial position towards the platform) and transit time (time taken to traverse the rod from the far end to the platform).

### Anle138b treatment

Mice were treated from 9 to 12 months of age with anle138b, starting at a time when loss of striatal DA release was already present in MI2 mice but prior to significant nigral neuron loss and completed when MI2 mice presented with no more than 30% cell loss. Drug or placebo were administered in standard diet (2g/kg of food, ssniff Spezialdiäten GMbH) throughout the whole 3 month study period to different cohorts of MI2 and C57Bl/6S mice. The levels of anle138b in plasma and brains of mice following this type of treatment have been previously reported (Wagner et al., 2015). At 12 months of age, mice were assessed using *in vivo* microdialysis. Subsequently brains were collected for analysis, as described above.

### Statistical analysis

The results are expressed as means ± SEM. A t-test was used for comparisons between two experimental groups. When more than two means were compared, one-way or two-way ANOVA, followed by a Bonferroni correction for test multiple comparison was used. Data from the *in vivo* microdialysis experiments were analyzed using two-way or three-way mixed ANOVA, with sampling time point as within-subject factor.

## RESULTS

### Generation of 1-120hαSyn MI2 mouse line

MI2 mice express truncated 1-120hαSyn under the control of the rat TH promoter (Figure 1A), but no endogenous mouse αSyn due to a spontaneous deletion of the *SNCA* gene in the C57Bl/6OlaHsd (C57Bl/6S) mice (Specht and Schoepfer, 2001; Tofaris et al., 2006). A C-terminally truncated form of αSyn was used because it has been shown to aggregate faster than the full-length protein (Crowther et al., 1998) and its presence has been shown in brain extracts from Parkinson’s and dementia with Lewy bodies patients (Baba et al., 1998). Immunoblotting revealed the presence of the transgenic protein in brain regions and neurons with TH-expression, such as substantia nigra (SN), striatum and olfactory bulb (OB), while no transgenic protein was expressed in cortex or cerebellum, where TH expression is not prominent (Figure 1B). Expression of transgenic 1-120hαSyn appeared significantly lower than that of endogenous αSyn in wild-type (wt) C57Bl/6J mice (Figure 1C) possibly because αSyn is present in different neuronal populations in wt mice, while its expression in MI2 mice is limited to TH^+^ neurons.

By immunohistochemistry, the distribution of 1-120hαSyn also followed the TH expression pattern, staining being evident in SN pars compacta (SNpc) and ventral tegmental area neurons, while no staining was evident in the SN pars reticulata, where αSyn is instead found in wt C57Bl/6J mice (Figure 1D). In the SNpc 1-120hαSyn expression was localized in both neuronal cell bodies and processes, while in the striatum, where nigral neurons project, it appeared as diffuse punctate staining (Figure 1D).

### Progressive aggregation of transgenic 1-120hαSyn

Immunohistochemical analysis of 1-120hαSyn in the nigrostriatal system of MI2 mice showed an age-dependent progressive accumulation of the protein. At 1.5 months, anti-αSyn antibodies stained small puncta that were diffusely distributed throughout the cell body of nigral neurons as well as numerous swollen processes. A more intense pattern was observed at 6 months, and at 12 months of age when besides the punctate αSyn immunoreactivity, large LB-like structures with condensed αSyn staining were seen, and neuronal processes were uniformly stained, with some containing large 1-120hαSyn inclusions (Figure 2A). None of these features were present in wt C57Bl/6J mice, where at 12 months of age, normal αSyn puncta were much smaller and less frequent than the inclusions in the MI2 mice (Figure 2A).

In the striatum the distribution of 1-120hαSyn was consistent with its synaptic enrichment, confirmed by co-staining with synaptobrevin (VAMP2), (Figure S1A) and followed the pattern of nigrostriatal projections. Staining for transgenic human 1-120hαSyn appeared as small puncta, accompanied by larger inclusions, whose number increased with age (Figure 2B), while wild type C57Bl/6J mice displayed a normal diffuse and homogeneous αSyn punctate pattern (Figure 2B). No αSyn staining was present in wt C57Bl/6S mice with deletion of the endogenous αSyn gene. Staining for VAMP2 showed a progressive redistribution of the protein at the synapse (Figure S1B), a pattern similar to that previously reported in our α-Syn120 mice showing SNARE protein redistribution (Garcia-Reitbock et al., 2010).

Immunoblotting of SN extracts showed a significant reduction of monomeric 1-120hαSyn in 12 month-old MI2 compared with 6 and 1.5 month-old MI2 mice with an associated increase in higher molecular weight (HMW) 1-120hαSyn species (Figure 2C). In the striatum, monomeric 1-120hαSyn also increased between 1.5 and 6 months of age and an increase in HMW species was present at 12 months of age, similar to the SN (Figure 2C). No apparent difference in monomeric 1-120hαSyn was found in the OB between 1.5-, 6- and 12 month of age, although an increase in HMW bands was also present in this region (Figure 2C).

### Super-resolution imaging of striatal 1-120hαSyn

In order to obtain a more detailed analysis of synaptic 1-120hαSyn at a near single-molecule resolution, we imaged the transgenic protein in the striatum of mice at 1.5, 6 and 12 months of age using *d*STORM following immunofluorescence staining with a human αSyn specific antibody (Figure 3A). To perform quantitative analysis of the aggregation profile, size, shape, inner density, local density per region of interest and percentage of aggregated 1-120hαSyn, a semi-automated software was built that implements a novel set of meta-analysis clustering algorithmic tools for analyzing super-resolution data (Figure S2). Using this approach, we observed progressive aggregation of 1-120hαSyn and followed the size distribution of the aggregates over time in the striatum of 1.5, 6 and 12 month-old mice. The population of aggregates was divided into 4 groups according to size: small, with a diameter ranging from 20-100 nm, medium, with a diameter between 100-300 nm, large, with a diameter of 300-500 nm and very large, with a diameter above 500 nm. *d*STORM imaging and analysis demonstrated an age-dependent increase in the overall number of 1-120hαSyn aggregates from 1.5 to 6 and 12 months of age(Figure 3B). This increase was mainly due to an age-dependent increase in small (20-100nm), medium (100-300nm) and large (300-500nm) aggregates. The number of very large aggregates (>500 nm) did not significantly change with age (Figure 3C).

**Figure 3.**
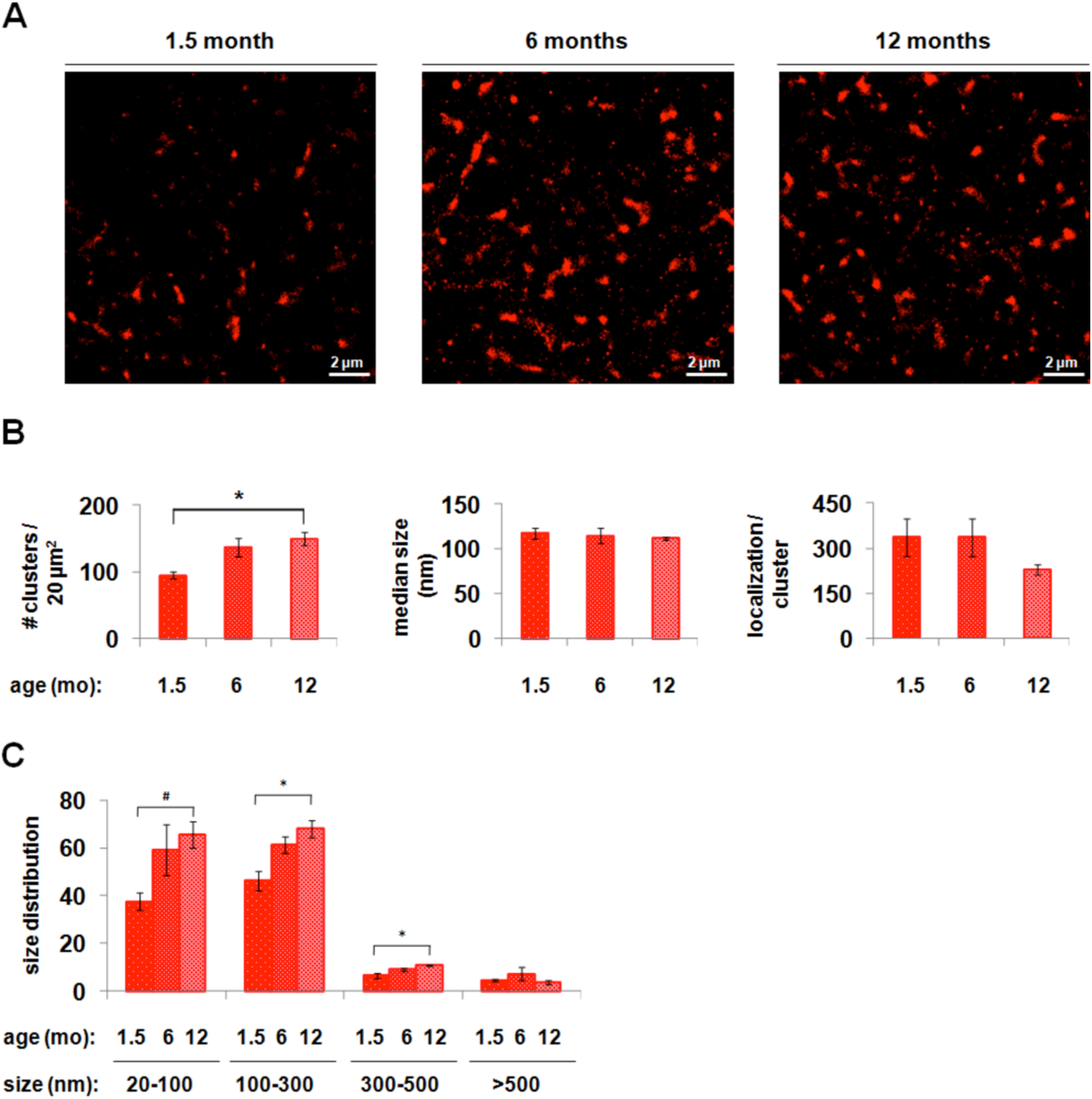
*d*STORM analysis of progression of 1-120 hαSyn aggregation in the striatum of MI2 mice. (A) Representative *d*STORM images of 1-120 hαSyn staining in striatum of MI2 mice at 1.5, 6 and 12 months of age showing increasing number of aggregates with age. (B) Quantification of *d*STORM data (see also Figure S2) (mean ± SEM, n=3 mice, one-way ANOVA, multiple comparisons with Bonferroni correction). The main effect of age is the increasing number of aggregates (clusters) (*p<0.05). No differences in the aggregate median size or number of localizations per cluster (e.g. the inner density which measures the number of fluorescent flashes in a cluster) were found. (C) Analysis of cluster size distribution shows a statistically significant increase in the number of small-size aggregates (20-100 nm, t-test #p<0.05), medium size aggregates (100-300 nm, one way ANOVA, *p<0.05) and large-size aggregates (300-500 nm, one way ANOVA, *p<0.05), between 1.5 and 12 months of age (See detailed statistical evaluation in Suppl. Material)

### Dopaminergic dysfunction in the striatum of MI2 mice

To establish whether the progressive aggregation of 1-120hαSyn affected DA neurotransmission in the striatum in MI2 mice, the total content of striatal DA was measured at 3, 6 and 12 months of age. No difference was detected between MI2 and C57Bl/6S control mice at 3 and 6 months of age, while a significant reduction in total DA levels was present at 12 months (control, 65.3±5.8, MI2, 38.1±5.7 pmol DA/mg of tissue, Figure 4A). DA content in 12 month-old MI2 mice was also reduced compared to MI2 mice at 3 and 6 months of age (87±9.2 and 73.1±11.1 pmol DA/mg of tissue, respectively; Figure 4A). No difference was found in DA levels at 12 months of age between control mouse lines with (C57Bl/6J) and without (C57Bl/6S) endogenous αSyn (Figure S3A), confirming that the absence of endogenous αSyn does not affect striatal DA levels, as previously shown (Garcia-Reitboeck et al., 2013).

**Figure 4.**
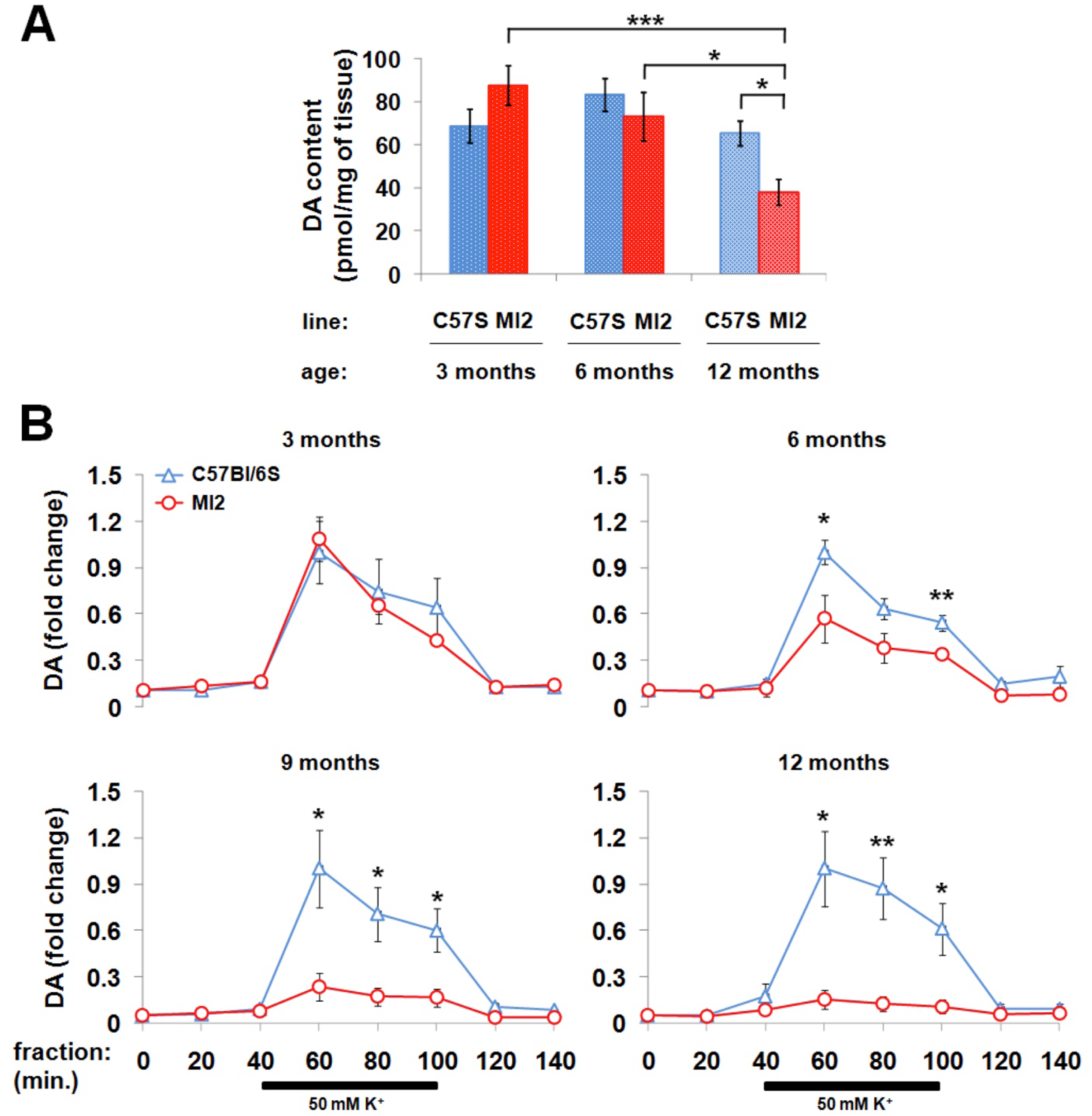
Striatal dopaminergic deficit in MI2 mice. (A) DA was measured in striatal lysates of MI2 and C57Bl/6S mice at 3, 6 and 12 months of age (mean ± SEM, n=5-8 mice per group). A main effect of age and interaction between genotype and age was identified by two-way ANOVA with Bonferroni correction, showing a significant reduction of DA levels in MI2 mice at 12 months compared to 12 months old C57Bl/6S mice, and to 3 month old and 6 month old MI2 mice (*p<0.05, ***p<0.001) (see also Figure S3A). (B) Striatal DA release was measured by *in vivo* microdialysis following the infusion of 50 mM KCl for 60 min between 40 to 100 min of the experiment. Data are expressed as a fold difference compared to the baseline fraction (0 min), normalized to the value obtained in age-matched C57Bl/6S controls at 60 min (mean ± SEM, n=4-6 mice). At 3 months of age no difference between MI2 and C57Bl/6S mice was observed, but a significant progressive decrease in DA release was found at 6, 9 and 12 months of age in MI2 compared with C57Bl/6S mice (see also Figure S3B). Two-way mixed ANOVA revealed a significant interaction between genotype and sample time at 6, 9 and 12 months of age (*p<0.05, **p<0.01, t-test for individual sampling time points). (See detailed statistical evaluation in Suppl. Material).

Dopaminergic deficit in MI2 mice was further investigated by measuring striatal DA release using *in vivo* microdialysis. DA release was induced in freely moving mice by infusion of 50 mM K^+^. No difference in DA release was observed between MI2 and control C57Bl/6S mice at 3 months of age, while at 6 months, a significant reduction in K^+^ stimulated DA release was present in MI2 mice compared with C57Bl/6S controls (samples were measured at 60, 80 and 100 min, normalized to peak C57Bl/6S value (at 60 min)) (0.57±0.15 vs 1±0.08 at 60 min, 0.38±0.09 vs 0.63±0.07 at 80 min, and 0.34±0.03 vs 0.54±0.05 at 100 min in MI2 vs control mice; Figure 4B). This impairment was progressive, with the loss of induced DA release being more prominent at 9 months of age (0.24±0.09 vs 1±0.25 at 60 min, 0.17±0.06 vs 0.71±0.17 at 80 min, and 0.17±0.06 vs 0.6±0.14 at 100 min in MI2 vs control mice) and further exacerbated at 12 months of age (0.16±0.06 vs 1±0.24 at 60 min, 0.13±0.05 vs 0.87±0.2 at 80 min and 0.11±0.04 vs 0.61±0.17 at 100 min in MI2 vs control mice; Figure 4B). No difference was observed in striatal DA release between C57Bl/6S and C57Bl/6J control mice at 12 months of age (Figure S3B). These findings indicate that striatal DA release impairment in MI2 mice is associated with the progressive aggregation of 1-120hαSyn. Importantly, a significant reduction in DA release in MI2 mice appears earlier than a reduction of total DA content in striatal tissue extracts, supporting the presence of a specific synaptic dysfunction (Figure 4).

### Loss of dopaminergic neurons in SNpc of MI2 mice

Nigral DA neuron death in the MI2 mice was investigated by unbiased stereological counting of cells stained for TH. The results showed a progressive TH^+^ cell loss in SNpc of MI2 mice compared to C57Bl/6S mice starting at 9 months of age (MI2, 11174±825.1 vs control, 13222±1658.8 TH^+^ neurons) becoming significant at 12 months of age (31% reduction in MI2 vs control mice: 8530.4±751.9 (MI2) vs 12398.8±946.1 (C57Bl/6S)) and further progressed at 20 months of age (54% reduction in MI2 vs controls: 6345.3±523.1 (MI2) vs 13836±616.1 (C57Bl/6S) TH^+^ neurons, respectively) (Figure 5A, B). The number of TH^+^ cells in SNpc of 20 month-old MI2 mice was also significantly lower (by 43%) compared to that in 9 months-old MI2 mice (Figure 5B).

**Figure 5.**
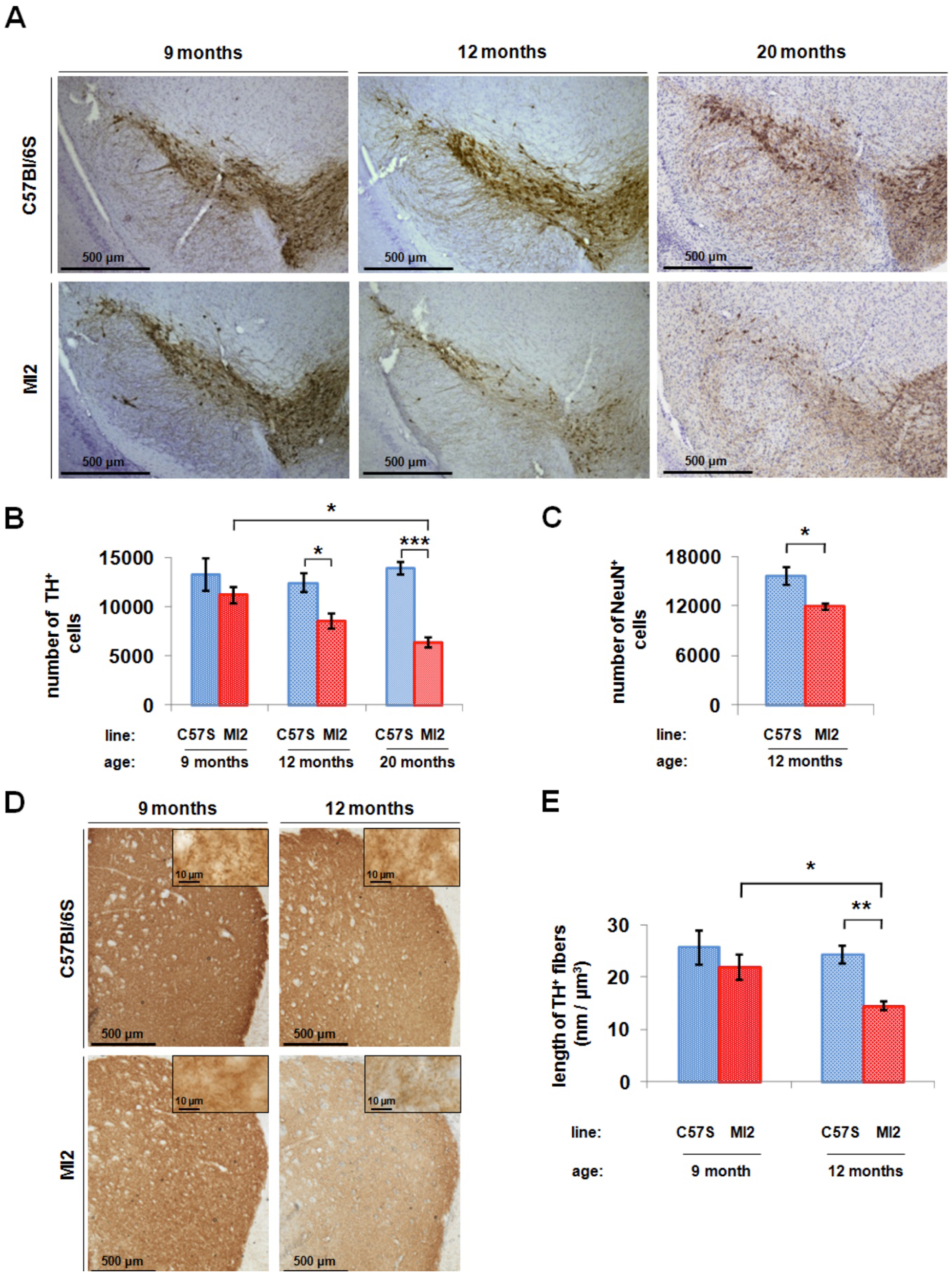
Loss of dopaminergic neurons in SNpc of MI2 mice. (A) Immunohistochemistry showed a reduction in TH staining in SNpc of MI2 mice compared to C57Bl/6S animals at 12 and 20 months of age. (B) Stereological counting of TH-positive neurons in SNpc. Average number of nigral TH-positive cells ± SEM, n=4-6 mice per group; two-way ANOVA with Bonferroni correction showing a significant reduction of TH^+^ neurons in MI2 mice compared to C57Bl/6S controls at 12 (*p<0.05) and 20 (***p<0.001) months of age. There was also a significant reduction in TH^+^ in MI2 mice between 9 and 20 months. (C) Stereological counting of total, NeuN^+^ neuron number in SNpc at 12 month of age (see also Figure S4). A significant decrease of NeuN^+^ neurons is present in MI2 mice compared to C57Bl/6S mice (average number of nigral NeuN-positive cells ± SEM, n=4 mice per group; t-test, *p<0.05). (D) Reduced TH-positive fiber staining in striatum of MI2 mice compared to C57Bl/6S mice at 12 months of age. Insets show high magnification images. (E) TH^+^ striatal neurites were estimated using the spaceball probe. (Mean length of striatal TH^+^ fibers per volume of striatal tissue ± SEM, n=4-6 mice per group; effect of genotype was identified by two-way ANOVA). Significant decrease in total length of TH^+^ fibers was present in 12 month-old MI2 mice compared to 12 month-old C57Bl/6S mice (**p<0.01) and 12 and 9 month old MI2 animals (*p<0.05), multiple comparisons with Bonferroni corrections. (See detailed statistical evaluation for 5B, 5C, 5E in Suppl. Material).

To confirm neuronal loss in SNpc and rule out the possibility that the decrease in TH cell number was reflecting a possible reduction in TH expression, cell counting was performed in 12 month-old mice following staining with the neuronal marker NeuN. NeuN and TH staining images were overlapped to delimit the counting area to the SNpc (Figure S4). The results showed a 20% reduction of NeuN positive neurons in SNpc of 12 month-old MI2 mice compared to C57Bl/6S (11877±383.5 (MI2) and 15633.5±1020.7 (C57Bl/6S) cells; Figure 5C). The reason for the reduced proportional neuronal loss observed following NeuN staining compared to TH staining is due to the fact that NeuN recognizes both TH and non-TH neurons in the SNpc, indeed, the absolute number of cell loss was similar for TH and NeuN stained neurons (−3868 and −3757 neurons, respectively).

We then assessed changes in DA innervation in the striatum of MI2 mice at different ages. We observed a significant decrease in striatal TH ^+^ innervation at 12 but not at 9 months in MI2 mice compared to C57Bl/6S mice (Figure 5D). In order to determine the loss of striatal innervation, we evaluated the density of TH-positive fibers by estimating their length in a defined volume of striatal tissue. We found that MI2 mice had a 40% loss of striatal TH innervation compared to C57Bl/6S mice (14.49±0.77 and 24.29±1.76 nm per µm^3^, respectively) at 12 months of age while no significant difference was present in 9 month-old MI2 mice compared to control (21.88±2.48 and 25.6±3.31 nm per µm^3^, respectively; Figure 5E). These results indicate that similar to the loss of nigral DA neurons, loss of nigrostriatal DA innervation occurs after the functional synaptic deficit, reflected by a reduction in DA stimulated release that starts at 6 months of age.

### Motor impairment in MI2 mice

An accelerating rotarod test was used to analyze motor function in MI2 mice at 6, 12, 15 and 20 months of age. An age-related gradual decline in rotarod performance was present in both C57Bl/6S and MI2 mice, but the reduction was more pronounced in MI2 mice, resulting in a significant loss of latency to fall from the rotarod at 20 months of age compared to their performance at 6 and 12 months of age (127.8 ± 14.9 s, 20m, 215.8 ± 9.9 s, 12m, 220.7 ± 11.5 s, 6m; Figure 6A). The performance of MI2 mice at 20 months of age was also significantly impaired compared to age-matched C57Bl/6S controls (127.8 ± 14.9 s and 181.7 ± 21.6 s, respectively; Figure 6A).

**Figure 6.**
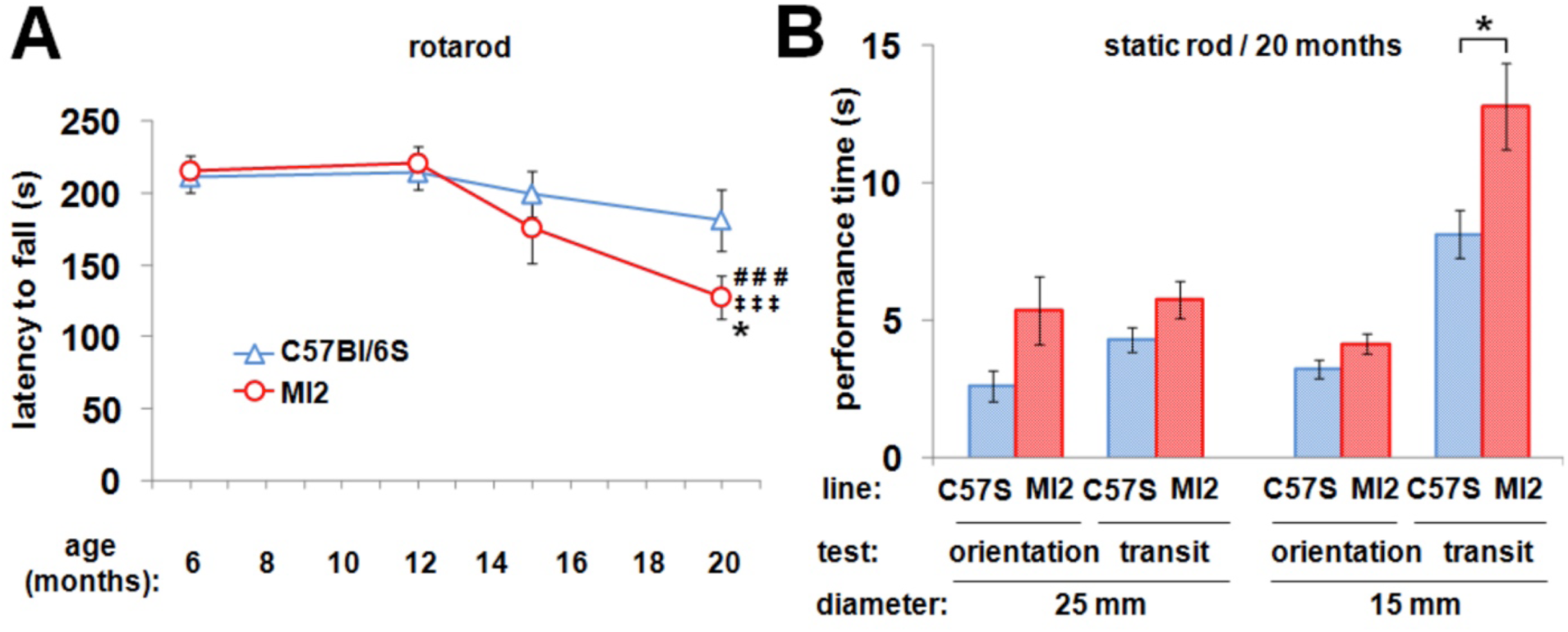
Motor impairment in aged MI2 mice. (A) C57Bl/6S and MI2 mice were analyzed using accelerating rotarod test at 6, 12, 15 and 20 months of age (mean latency to fall from the rod ± SEM, n=12-18 mice per group). A main effect of age was identified by two-way ANOVA. Statistically significant differences between 6 and 20 months (^###^p<0.001) and 12 and 20 months (^‡‡‡^p<0.001) in MI2 mice, and between MI2 and C57Bl/6S animals at 20 months (*p<0.05) were revealed by multiple comparisons with Bonferroni correction. (B) Motor performance of 20 month-old mice was tested using the static rod test. (Mean orientation time or mean transit time ± SEM, n=13-18 mice). No differences were identified between experimental groups using a 25 mm rod or in orientation time using a 15 mm rod, but there was a statistically significant difference in transit time on the 15 mm rod between MI2 and C57Bl/6S mice (*p<0.05, t-test).

The presence of a motor deficit was also tested in 20 month-old MI2 and C57Bl/6S mice using the static rod test where the time required to turn 180° from the initial position (orientation time) and the time required to travel from the far end of the rod to the platform (transit time) were measured. We found no significant differences between transgenic and control mice in either orientation or transit time when a 25 mm diameter rod was used, but when the test was performed with a 15 mm rod, although there was no significant difference in the orientation time, the transit time was longer for MI2 mice compared to controls (12.8±1.6 s and 8.1±0.8 s, respectively), confirming a selective motor impairment in MI2 mice (Figure 6B).

### The oligomer modulator, anle138b, rescues DA deficit in MI2 mice

MI2 mice reproduce key features of PD, progressive aggregation of αSyn in the nigrostriatal system, progressive impairment of striatal DA release, reduction in total striatal DA, loss of nigrostriatal DA fibers and nigral DA neurons and motor impairment when the SNpc DA neuron loss reaches 50%. This line therefore appears appropriate for testing potential therapies for PD. Hence, we investigated if a small molecule modulator of aggregation, anle138b was able to rescue DA impairment in MI2 mice. Furthermore, we used *d*STORM imaging and analysis to determine how anle138b acts *in vivo* and how it affects the spatial distribution of striatal 1-120hαSyn.

Anle138b was administered in the food (2 g/kg of food) for 3 months to MI2 and C57Bl/6S mice. Treatment with anle138b or placebo started at 9 months of age, when striatal DA release impairment is already present in MI2 mice, and ended at 12 months of age, when 30% nigral neuron loss is detected in non-treated animals (Figure 7A). Using immunohistochemistry, we found that anle138b treatment reduced 1-120hαSyn aggregation in both SNpc cell bodies and synaptic terminals in the striatum (Figure 7B). Application of *d*STORM analysis not only confirmed the reduction in striatal 1-120hαSyn aggregation but also provided a mechanistic explanation for anle138b activity. *dS*TORM analysis revealed two opposite but complementary effects of anle138b: we detected an increased percentage of small, non-clustered 1-120hαSyn species (anle138b, 53.4±2.9%; placebo, 29.9±3%; Figure 7C, D) that occurred concomitantly with a reduction in the inner density of small and large size aggregates (anle138b, 140.9±12.7; placebo, 233.5±25.3 fluorophore localizations; Figure 7C, D). The total number of aggregates was only slightly but not significantly reduced following anle138b treatment (Figure 7D). Analyses of striatal extracts by immunoblotting confirmed this result showing a reduction in HMW 1-120hαSyn band and an increase in monomeric protein (Figure 7E). These data show that anle138b substantially affects the aggregation profile of 1-120hαSyn in MI2 mice.

Accordingly, we speculated that the non-aggregated small 1-120hαSyn species, revealed by *d*STORM, whose number increased following anle138b treatment may represent monomers or small 1-120hαSyn assemblies. To examine this hypothesis, we compared the size distribution of the non-aggregated population in MI2 mouse striatum to the size distribution of recombinant monomeric hαSyn. Recombinant hαSyn was plated at low density onto a coverslip, immunolabeled, and imaged using the same protocol as that used to visualize 1-120hαSyn in MI2 striatal slices. Mapping the recombinant hαSyn population resulted in a size distribution with a mean diameter of 23 nm and an average of 7 localizations per protein (Figure S5A,B). Using the same parameters of clustering, we observed that following anle138b treatment, 85% of 1-120hαSyn population in MI2 mouse striatum had a very similar size distribution with a mean diameter of 23 nm and an average of 7 localizations per protein (Figure S5C,D). These data suggest that anle138b treatment resulted in an increase most probably of monomeric and small assemblies of monomeric hαSyn species that could have been possibly either released from dissolving aggregates or not recruited to them.

To determine the effect of anle138b on dopaminergic impairment, we measured striatal DA release in MI2 mice. A 3-month treatment was started at the age of 9 months, when the DA release deficit was already present. Released DA was significantly increased in anle138b-treated MI2 mice compared to placebo-treated MI2 animals (K^+^-stimulated DA release, relative values of DA release in anle138b-vs placebo-treated MI2 mice sampled at 60, 80 and 100 min, after normalized to peak C57Bl/6S mouse fraction value (at 60 min) reached 0.69±0.14 vs 0.13±0.02 (60 min), 0.36±0.07 vs 0.07±0.01 (80 min) and 0.28±0.05 vs 0.09±0.02 (100 min)). The DA release values in C57Bl/6S anle138b-vs placebo treated mice were not altered (1.03±0.25 vs 1±0.12 (60 min), 0.41±0.09 vs 0.55±0.05 (80 min) and 0.38±0.09 vs 0.39±0.03 (100 min) (Figure 8A). We then analyzed the effect of anle138b treatment on the number of nigral DA neurons and found a significant increase in number of TH^+^ neurons in SNpc of anle138b-treated MI2 animals compared to placebo-treated mice (Figure 8B), with anle138b preventing cell loss. The number of nigral TH^+^ cells did not differ between C57Bl/6S mice treated with either placebo or anle138b (10886.7±706.5 and 10792±858.9 TH^+^ cells, respectively) and anle138b-treated MI2 mice (9944±567.5 TH^+^ cells), while a substantial DA neuron loss was present in placebo-treated MI2 mice (7465±41.4 TH^+^ cells) (Figure 8C). Taken together, these results show that targeting αSyn aggregation by anle138b rescues the DA release deficit, and protects against cell loss, even when treatment was started when synaptic functional impairment was already present.

**Figure 7.**
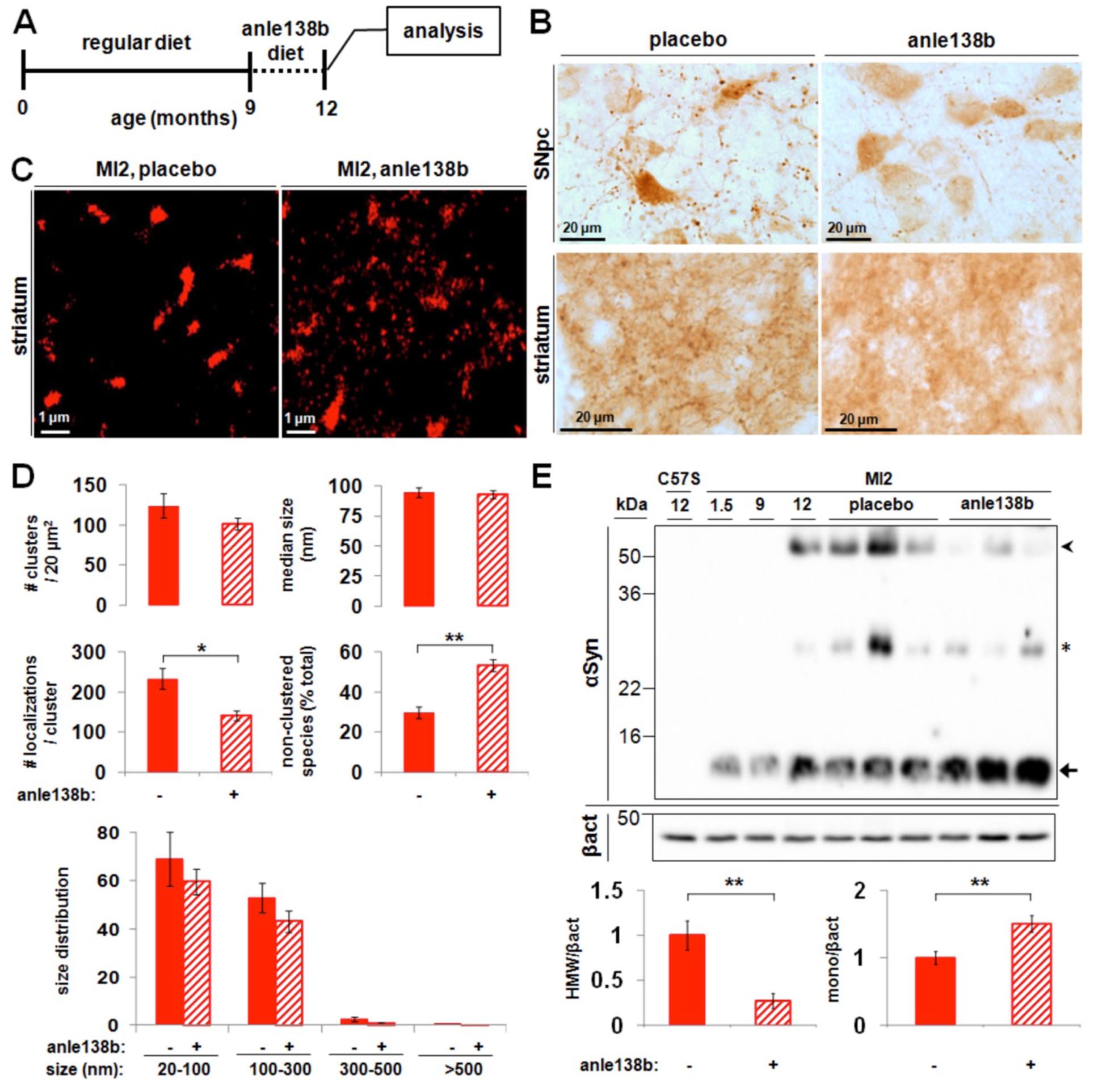
Effect of anle138b on αSyn aggregation in MI2 mice. (A) Mice were treated with anle138b starting at 9 months and analyzed at 12 months of age. (B) Treatment with anle138b reduces the accumulation of 1-120hαSyn in SNpc and striatum of MI2 mice as shown by Syn1 antibody immunostaining. (C) Representative *d*STORM images of 1-120 hαSyn staining in striatum of 12 months old MI2 mice treated with placebo (left panel) or anle138b (right panel). Large aggregates appear less dense and an increased number of smaller species are seen following anle138b treatment. (D) Quantification of *d*STORM data revealed a significant decrease in the inner density of aggregates (#localizations/cluster) and a significant increase in the percentage of non-clustered 1-120hαSyn in anle138b-treated MI2 mice compared to placebo-treated littermates (see also Figure S5) (mean ± SEM, n=3 mice; t-test, *p<0.05, **p<0.01) while the number of clusters and their size did not change significantly. (E) Immunoblotting of striatal brain extracts from 12 month-old mice shows a decrease of αSyn high molecular weight (HMW) bands (arrowhead) and an increase of monomeric αSyn (arrow) in striatum of MI2 anle138b-treated mice compared to MI2 placebo treated. 1-120 hαSyn levels are also shown for 1.5, 9 and 12 month-old untreated MI2 mice. C57Bl/6S (C57S) not expressing endogenous αSyn are shown as negative control. The star denotes non-specific bands recognized by Syn1 antibody. Blot shows levels of αSyn before and after anle138b in 3 mice representative of 5-6 mice per group tested. Bottom panel - quantification of HMW 1-120hαSyn and monomeric 1-120hαSyn after normalization to the levels of β-actin in all 5-6 mice tested. (Fold difference compared to placebo-treated mice ± SEM, n=5-6 mice per group, t-test: **p<0.01).

**Figure 8.**
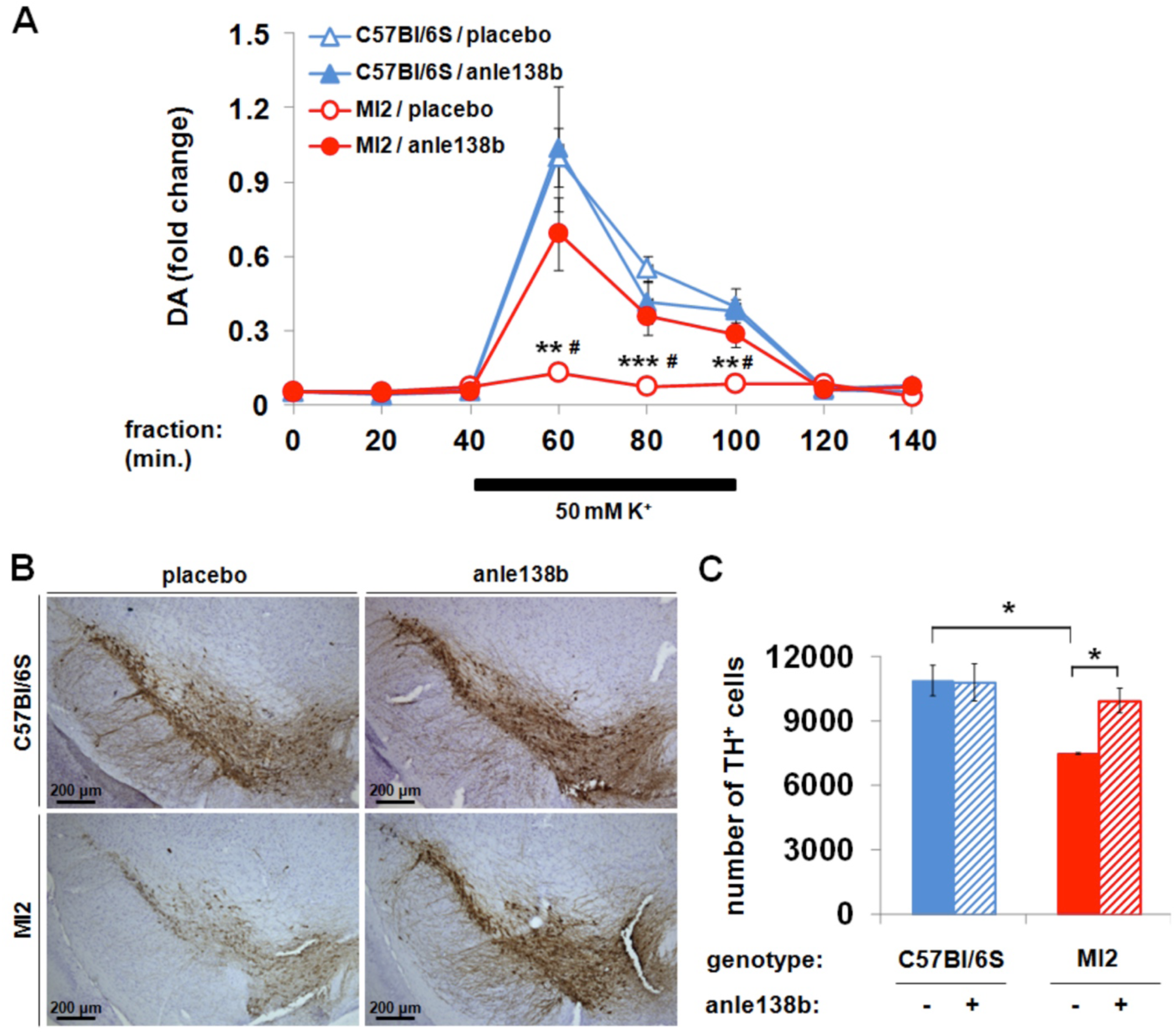
Effect of anle138b on striatal DA release and SNpc DA neuron death in MI2 mice. (A) The effect of anle138b treatment on striatal DA release studied by *in vivo* microdialysis. Data are normalized to the value obtained in the C57Bl/6S/placebo group at 60 min (means ± SEM, n=5-6 mice per group). Three-way mixed ANOVA identified an effect of genotype, sampling time, two-way genotype × sampling time interaction, and sampling time × genotype × treatment three-way interaction. Statistically significant differences in K^+^-induced DA release at 60, 80 and 100 min were found between placebo-treated MI2 and C57Bl/6S mice (**p<0.01, ***p<0.001) and between placebo- and anle138b-treated MI2 mice (^#^p<0.01) (multiple comparisons with Bonferroni correction). (B) Preservation of TH immunoreactivity in SNpc of anle138b-treated MI2 mice compared with placebo-treated littermates. (C) Effect of anle138b treatment on nigral DA neuronal death determined using stereology (average number of nigral TH-positive cells ± SEM, n=3 mice per group). A main effect of genotype was identified by two-way ANOVA. Anle138b treatment resulted in increased number of TH^+^ cells in MI2 mice compared to placebo-treated littermates (*p<0.05). Placebo-treated MI2 mice had less nigral TH^+^ cells than placebo-treated C57Bl/6S mice (*p<0.05, multiple comparisons with Bonferroni correction). (See detailed statistical evaluation in Suppl. Material).

## DISCUSSION

In PD, αSyn aggregation is associated with a reduction of total striatal DA, a decrease in striatal DA release and nerve cell loss in the SNpc which, when reaching around 50%, leads to motor impairment (Brooks et al., 1990; Kordower et al., 2013). Animal models recapitulating the progressing nature of αSyn pathology and DA dysfunction in PD are essential for testing new therapies. Here we describe MI2 mice, a new transgenic mouse model that expresses 1-120hαSyn and shows a high similarity to PD pathological features with progressive loss of DA and a deficit in striatal DA release, as well as αSyn aggregation in SNpc neurons and their striatal terminals, and dopaminergic neuronal death in the SNpc that is associated with impaired motor behavior when cell death reaches around 50%, similar to the human condition.

Dopaminergic pathology in MI2 mice was associated with changes in 1-120hαSyn protein distribution in SNpc and striatum accompanied by redistribution of SNARE proteins such as VAMP 2 (Figure S1) similar to what we have previously described in transgenic α-Syn120 mice and PD brains (Garcia-Reitbock et al., 2010) and was also reported in DLB cases (Kramer and Schulz-Schaeffer, 2007). In SNpc of MI2 mice 1-120hαSyn appeared in neuronal cell bodies as multiple small inclusions at 1.5 months of age, while large LB-like structures were present in numerous cells by 12 months of age. Neuronal processes in the SNpc contained large dotted inclusions at 1.5 months of age, while in older mice processes were filled with dense aggregates. This pattern indicates a progression in αSyn aggregation from small to large structures, similar to human α-synucleinopathies (Kuusisto et al., 2003). In the striatum of 1.5 month-old MI2 mice, 1-120hαSyn deposits were small, but widely distributed in the neuropil, and in older animals they were more frequent and of larger size. Both, in SNpc and striatum, the αSyn aggregates became more insoluble with time, as shown by the appearance of stable HMW bands in immunoblots of MI2 mouse brain extracts. To further characterize the progression of striatal synaptic 1-120hαSyn aggregation, we used *d*STORM, which allows detection of protein distribution at single molecule level and 20nm resolution. Although *d*STORM has been successfully employed to study synaptic proteins (Bar-On et al., 2011; Bar-On et al., 2012; Bielopolski et al., 2014; Fulterer et al., 2018; Tang et al., 2016) we believe that this is the first time that *d*STORM is used to analyze the progression and rescue of presynaptic αSyn pathology *in vivo* in brains of a PD mouse model. *d*STORM confirmed a progressive aggregation of 1-120hαSyn in the MI2 mouse striatum. Of all the parameters analyzed, cluster abundance and density (number of clusters per 20 µm^2^) of 1-120hαSyn aggregates appeared to be the factors best associated with the reduction in striatal DA release. The presence of αSyn aggregates has been recently demonstrated by some of us using a different super resolution microscopy method (DNA PAINT) in induced pluripotent stem cells (iPSCs) from a patient with a triplication of the SNCA gene (Whiten et al., 2018), however, one of the unique finding of the present study is the differential age-dependent effect on aggregate size. Only 20 to 500 nm aggregates were the populations that increased with age, and these populations of small αSyn species cannot be observed using conventional microscopy, establishing *d*STORM as a novel approach for the analysis of αSyn aggregation *in vivo* in mouse models.

Striatal impairment of DA release was observed in MI2 animals before measurable loss of DA cells in SNpc and DA innervation in the striatum (6 vs 12 months). This is consistent with our previously reported α-Syn120 mice, where a deficit in striatal DA release was present in the absence of DA neuron death in SNpc (Garcia-Reitbock et al., 2010). This observation supports the hypothesis that synapses play a central role in PD pathogenesis, being the primary site of dysfunction before loss of neurites and neurons occur, supporting the notion of retrograde degeneration (Calo et al., 2016; Decressac et al., 2012). Moreover, overt motor phenotype occurred when around 50% of SNpc DA neurons were lost similar to human PD (Brooks et al., 1990; Kordower et al., 2013).

We then used MI2 mice to test whether DA pathology might be rescued by targeting neuronal αSyn aggregation with anle138b, which has previously been shown to have beneficial effects by interacting specifically with structural epitopes of various protein aggregates and to rescue protein aggregation and related pathological features in models with αSyn, prion and tau aggregation (Martinez Hernandez et al., 2018; Wagner et al., 2015; Wagner et al., 2013). Unlike the MI2 mice, the αSyn transgenic models used in previous anle138b studies did not present with consistent neuronal nigrostriatal αSyn aggregation or αSyn-related progressive dopaminergic impairment with motor impairment related in some cases more to spinal cord pathology (Wagner et al., 2013) or αSyn pathology in oligodendrocytes (Heras-Garvin et al., 2018). Thus, our study is the first to investigate the effects of anle138b in a model with nigrostriatal neuronal DA alterations and neuronal αSyn aggregation as observed in PD. We show that a 3 month-treatment with anle138b (started at 9 months of age, when a deficit in striatal DA release but no significant neuronal loss is present) rescues DA dysfunction and prevents nerve cell death. This is particularly significant, because it shows the presence of a time window, when DA function can still be rescued and neuronal death prevented despite the presence of αSyn-related synaptic alterations. Anle138b significantly changed 1-120hαSyn aggregation in the striatum and SNpc of MI2 mice, as observed by immunohistochemistry and immunoblotting. However, more information on its mechanism of action *in vivo* was obtained by *d*STORM, which showed an increase in the percentage of small, non-clustered αSyn species in the striatum following anle138b administration. These species are likely to represent monomers and small assemblies of monomeric αSyn as determined by comparing their size to the size of recombinant hαSyn spread on a coverslip and immunostained as in the brain tissue and similarly examined with *d*STORM. This analysis was an approximation, since it compared the transgenic truncated 1-120 protein with full-length recombinant hαSyn but this was done because the recombinant truncated protein has a greatly increased tendency to aggregate (Crowther et al., 1998) making it difficult to determine the presence of *bona fide* monomeric species.

Moreover, after anle138b treatment, large aggregates showed a significant reduction in inner density. This suggests that anle138b *in vivo* could interfere with aggregation by preventing the formation of new aggregates and/or the recruitment of non-aggregated 1-120hαSyn into existing aggregates possibly destabilizing them, thus leading to a reduction of their inner density, resulting in both cases in an increase in small, non-clustered αSyn species in the environment. Whether the anle138b effect is related to one or both of these mechanisms could be established by monitoring aggregation of hαSyn with *d*STORM in anle138b-treated MI2 mice over time.

It remains to be clarified, how anle138b produced its beneficial effect, whether the effect is related to the reduced toxicity of less dense aggregates or to the increase in small αSyn species, which could be physiologically active and restore DA release. It will be interesting to see if removal of small αSyn species by inducing their degradation can potentiate the effects of anle138b by further reducing possible aggregation or reduce its effect, as it could be expected if the small αSyn species are functional.

In summary, we report here a transgenic mouse model that recapitulates the progressive αSyn aggregation and DA neuron dysfunction and death observed in human PD. Striatal synaptic DA dysfunction preceded SNpc DA neuronal death and loss of striatal TH terminals, supporting synaptic dysfunction as being an early pathogenic factor in PD. Anle138b restored DA function and prevented nerve cell loss by disrupting αSyn aggregate formation as shown for the first time by using *d*STORM. Overall, this study indicates that MI2 mice are a suitable model to test mechanism-based therapies for α-synucleinopathies, and *d*STORM is a useful technique for studying structural/morphological changes related to aggregate toxicity. This work also indicates that there is a window of time when it is possible to prevent DA neuronal death, even when striatal DA release is already impaired.

## ACKNOWLEDGEMENTS

We acknowledge the help of Ms Emma Carlson in the *in vivo* microdialysis experiments and preparation of tissue for immunohistochemistry. We thank Drs Aviva Tolkovsky and Michel Goedert for useful discussions and comments on the manuscript. We are also grateful to the personnel in the Bioscience facilities, in particular Ms Debbie Drage who has helped us generate and maintain the MI2 mouse line in that due to several changes of bioscience facilities the mice had to be re-derived and a new colony started several times over the years.

This work was supported over the years by Parkinson’s UK, the Cure PD Trust, the MJ Fox Foundation, UK Medical Research Council, Alzheimer’s Research UK, Neuroscience Network of Excellence (NNE), Israel Science Foundation grants 2546/16, and the Max Planck Society.

## AUTHOR CONTRIBUTION

M.G.S. planned and supervised the study and provided funding; A.G., C.G. provided anle138b; U.A. supervised the *d*STORM and contributed funding for it; M.W. performed the immunoblots, immunohistochemistry, microdialysis, stereological and behavioral experiments, maintained mouse colonies, and supervised the anle138b treatment; D.B-O. performed *d*STORM imaging and analysis; L.C. supervised treatment with anle138b, and assisted in microdialysis experiments; O.A. was involved in the initial characterization of the MI2 mice, and supervised the anle138b treatment; M.I. produced the transgene used to generate the MI2 line; J.X. and J.W.D. performed the dopamine measurements; S.R. and A.L. were involved in developing the anle138b compound; M.W. and M.G.S. wrote the manuscript with contributions from U.A., D. B-O. and C.G. The manuscript was revised by all the authors.

