## Supplementary Material for "Depopulation of α-synuclein aggregates is associated with rescue of dopamine neuron dysfunction and death in a new Parkinson’s disease model"

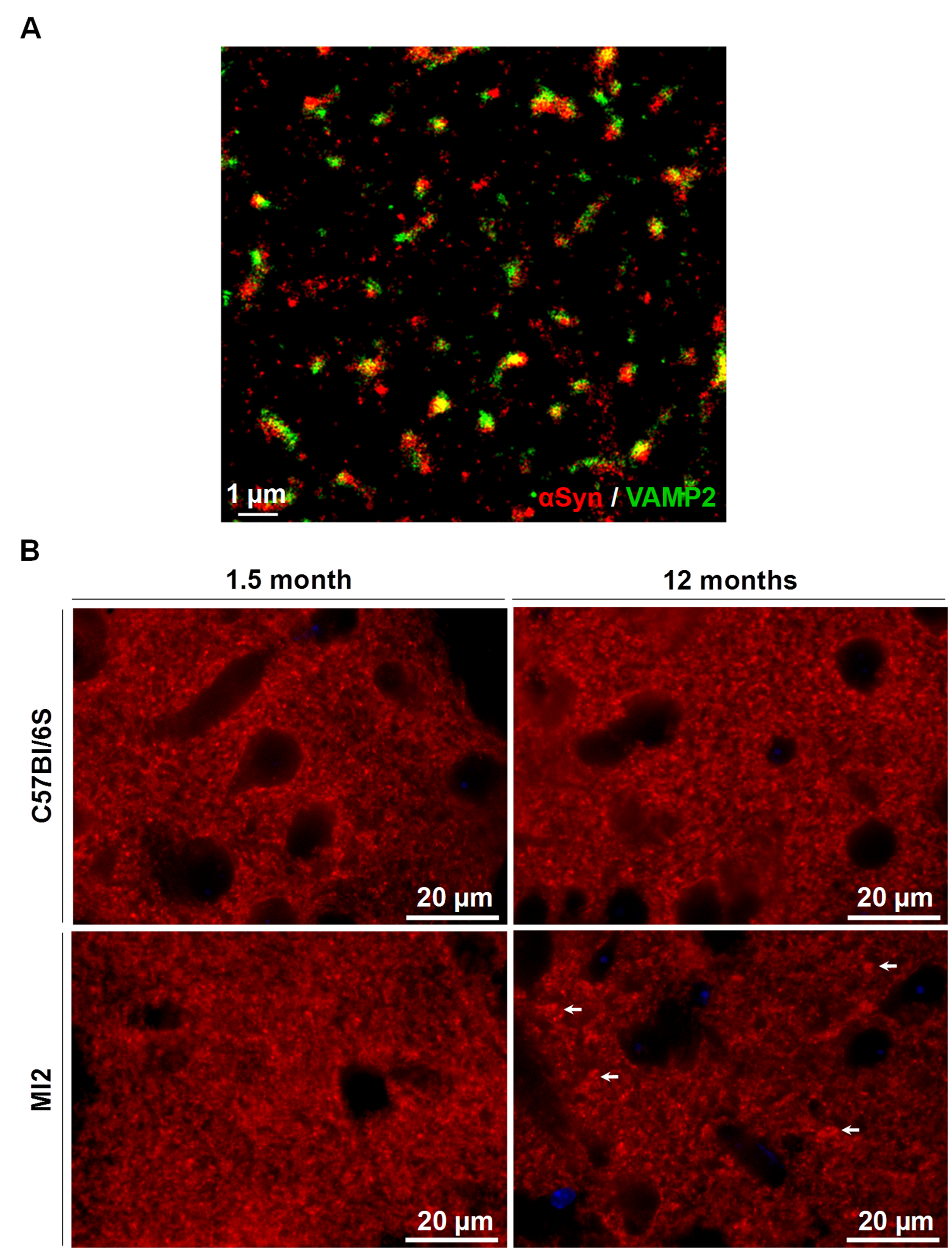

**Figure S1.** Related to Figure 1. Co-localization of 1-120hαSYN and VAMP2 in striatal pre-synaptic compartment and VAMP2 redistribution. (**A**) Localization of 1-120hαSYN in striatal synapses. Co-localization of 1-120hαSYN protein with synaptic marker, synaptobrevin (VAMP2), confirmed distribution of 1-120hαSYN protein in synaptic terminals in striatum of 6-months old MI2 mice. (**B**) Redistribution of VAMP2 in striatum of MI2 mice. Distribution of VAMP2 in control C57Bl/6S mice is homogenous at both 1.5 and 12 months of age. MI2 mice exhibit progressive redistribution of striatal VAMP2. At 1.5 month, VAMP2 is distributed homogenously, similar to C57Bl/6S mice, when at 12 months its distribution is visibly less homogenous, and numerous VAMP2-immunopositive clumps can be detected (arrows).

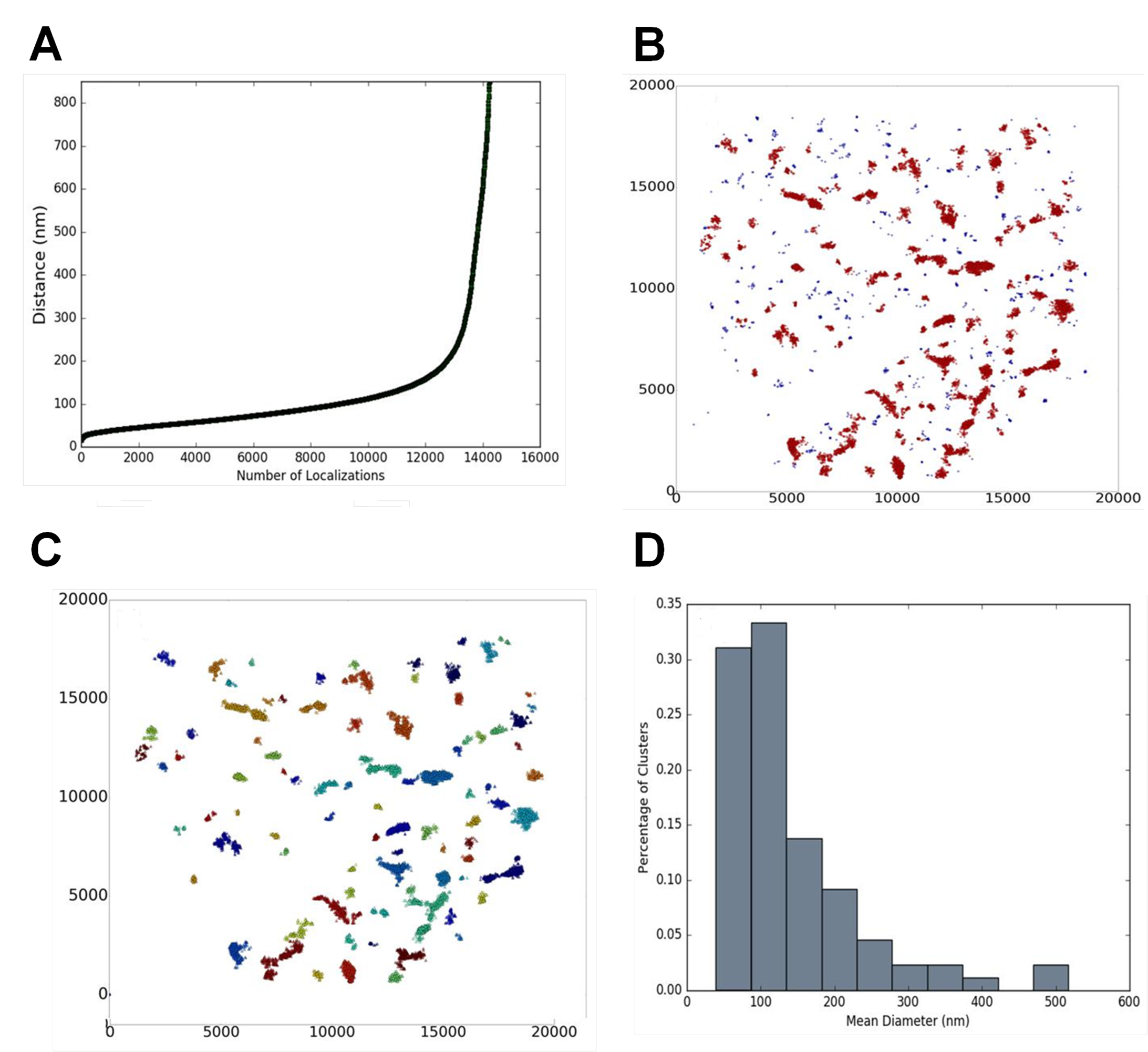

**Figure S2.** Related to Figure 3. Analysis of dSTORM data. (**A**) dbscan (density-based) cluster analysis was performed for the single localizations (predicted monomeric protein) in each image using the parameters of ε=200 and k=16 as determined by a set of k-dist measurement of nearest neighbors analysis (Bar-On et al., 2012). (**B**) Dividing protein localizations into clustered (aggregated; red) and non-clustered (non-aggregated / free (blue)) localizations (**C**) Different clusters / aggregates of αSYN are assigned with different colours, and for each aggregate the size, density, number of localization and shape is calculated. (**D**) Representative αSYN cluster size-distribution histogram in striatum of MI2 mice at 1.5 month of age.

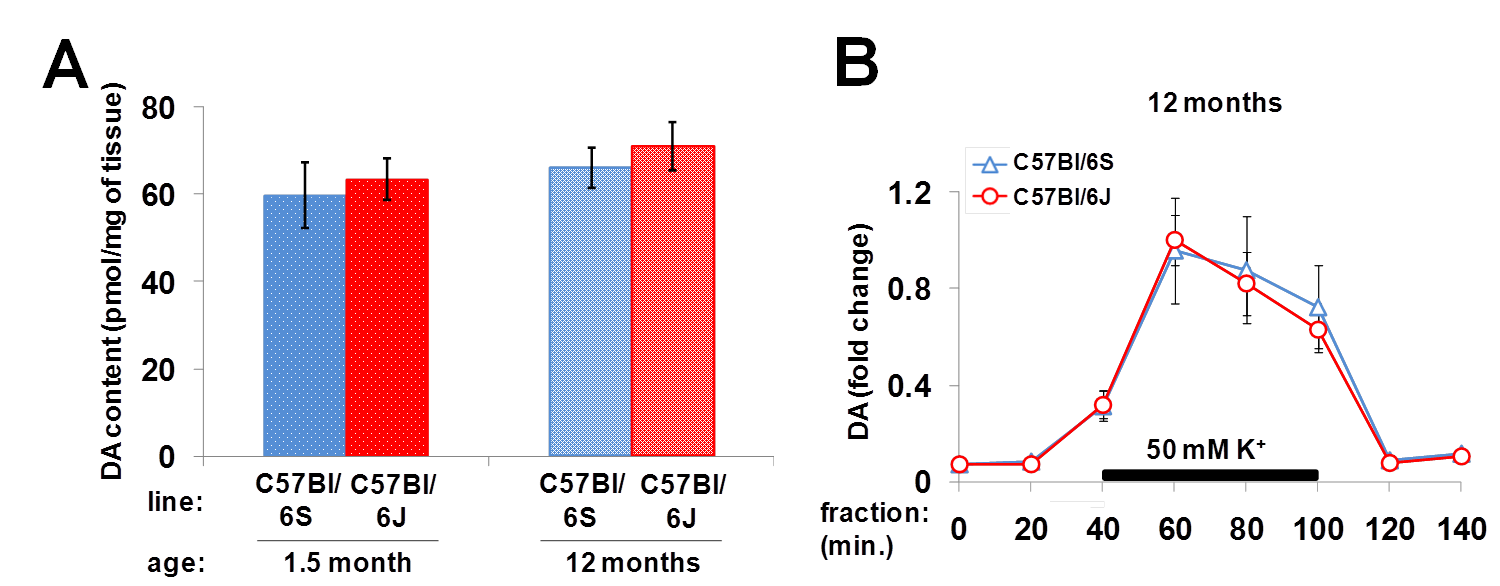

**Figure S3.** Related to Figure 4. Comparison of striatal dopamine in αSYN-null C57Bl/6S control mice and endogenous αSYN-positive C57Bl/6J mice. (**A**) Total content of DA in striatal tissue was unaltered between C57Bl/6S and C57Bl/6J mice at 1.5 and 12 months of age. (**B**) No difference in DA release, measured by microdialysis, was found in the striata of C57Bl/6S and C57Bl/6J mice at 12 months of age as we have also shown previously (Garcia-Reitbock et al., 2013).

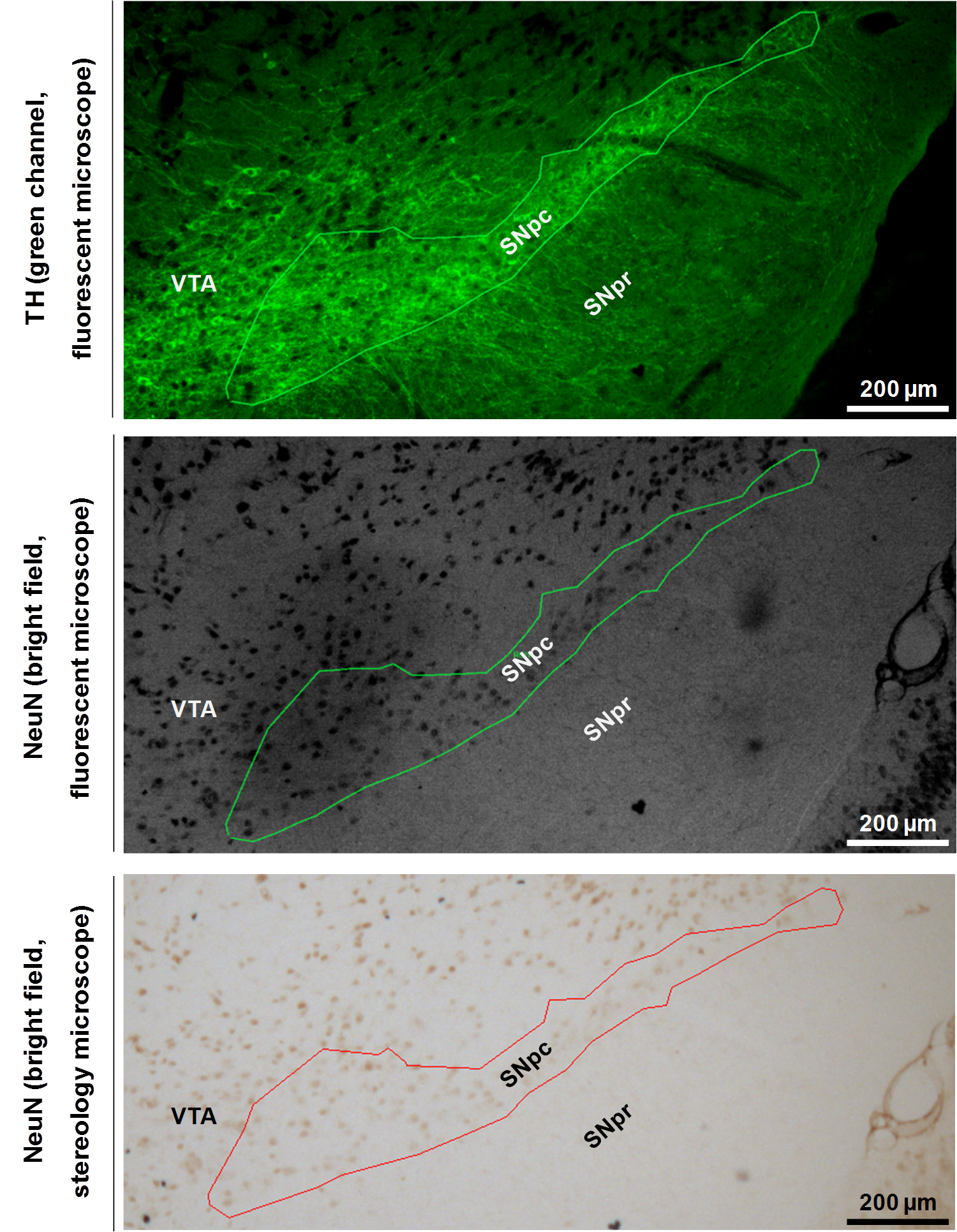

**Figure S4.** Related to Figure 5. Representative image of combined peroxidase-based immunostaining of NeuN and immunofluorescence staining of TH for stereological counting of nigral NeuN-positive neurons. The double-stained sections were first analyzed using epifluorescent microscope. SNpc was contoured according to TH immunofluorescence in the green channel (upper panel), and the contour was superposed on the bright field image of NeuN immunohistochemistry (middle panel). Slides were then transferred to a stereology microscope and the contour of SNpc was precisely reconstructed in the bright field image in Stereo Investigator (bottom panel) for stereological counting of NeuN-positive nigral neurons. SNpc = substantia nigra pars compacta, SNpr = substantia nigra pars reticulata, VTA = ventral tegmental area.

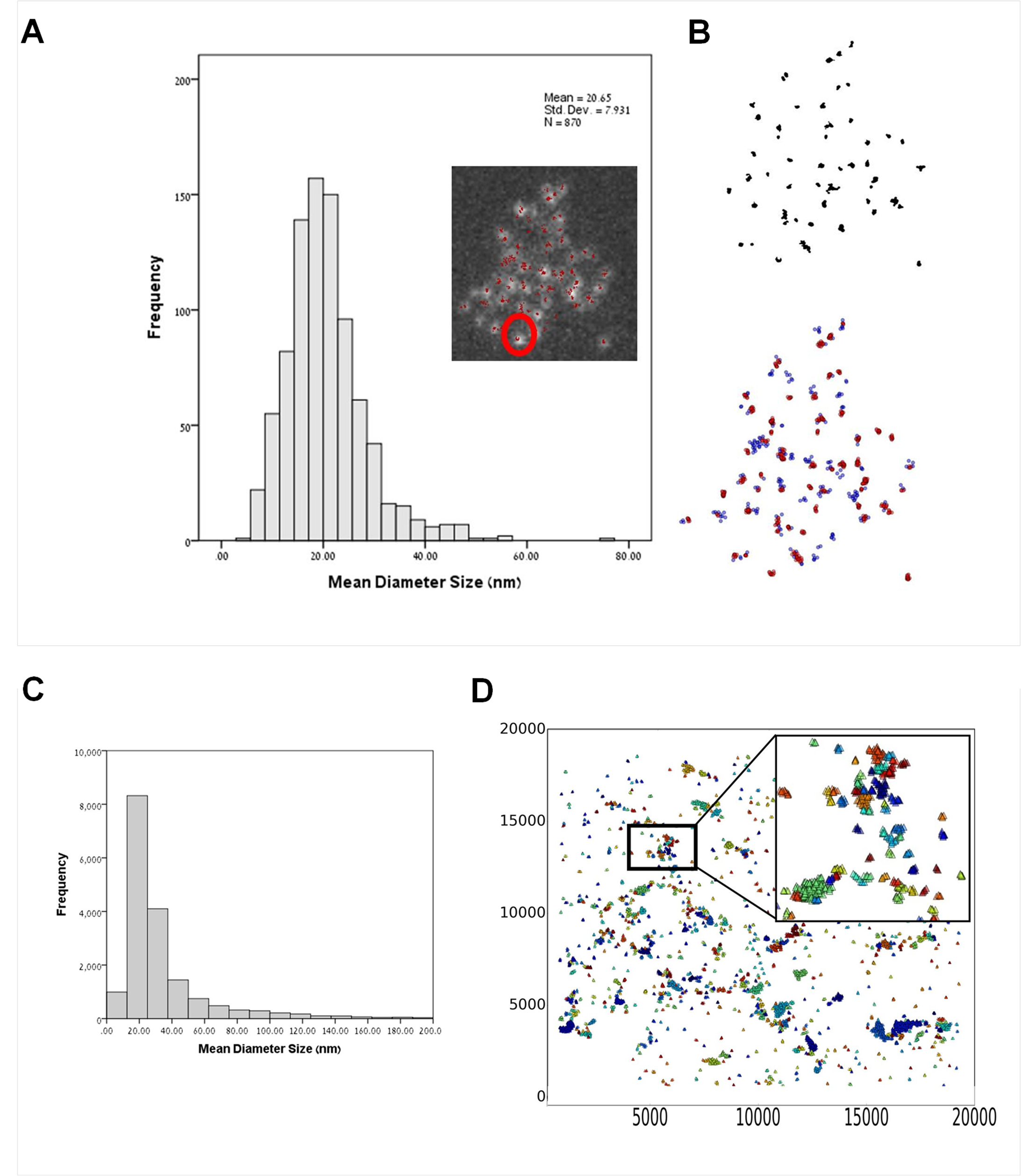

**Figure S5.** Related to Figure 7. Identification of monomeric 1-120hαSYN in striatum of anle138b-treated MI2 mice. (**A**) Size distribution for monomeric recombinant αSYN. Putative single αSYN protein was imaged by dSTORM and analysed using the dbscan analysis. The mean diameter size of a single αSYN was found to be 20.65 nm. Inset: wide field image of recombinant αSYN, with superposed fluorophore coordinates as imaged by dSTORM (indicated in red). An example of a putative single αSYN is marked by a red circle. (**B**) The parameters for a single αSYN protein were detected using a combination of imageJ analysis for finding centers of mass of proteins and dbscan analysis, both tools were combined to verify the definition of a single protein. (**C**) Size distribution of the non-clustered αSYN population in striatum of anle138b-treated MI2 mice. 85% of αSYN is in entities that would correspond to what we have defined as monomers (mean diameters size of 23 nm and median of ~7 fluorophores per cluster, similarly to the definition of the recombinant single protein. (**D**) A representative image of the non-clustered population composed of smaller low density dispersed aggregates and large population of monomers between them. Inset: representative dispersed aggregate, composed mainly of monomers (indicated in different colors).

**Detailed statistical evaluation of the results shown in the Figures**

**Figure 1B.** One-way ANOVA revealed a main effect on 1-120 αSYN normalized to either β-actin or TH (F(2,6)=99.813, p<0.001, and F(2,6)=22.97, p=0.002, respectively). Multiple comparisons with Bonferroni correction revealed statistically significant differences between SN and OB (p<0.001 for αSYN/β-act, and p=0.002 for αSYN/TH) and between Str and OB (p<0.001 for αSYN/β-act and p=0.005 for αSYN/TH).

**Figure 2C.** In SN, one-way ANOVA revealed a main effect of age on αSyn/β-actin (F(2,6)=10.540, p=0.011) and αSyn/TH (F(2,6)=6.416, p=0.032). Multiple comparisons with Bonferroni correction revealed statistically significant difference between 1.5 and 12 months (p=0.019) and between 6 and 12 months (p=0.026) for αSyn/β-actin, and between 6 and 12 months (p=0.042) for αSyn/TH. In striatum, one-way ANOVA revealed a main effect of age on αSyn/β-actin (F(2,6)=6.338, p=0.033) and αSyn/TH (F(2,6)=7.431, p=0.024). Significant differences were present for αSyn/β-actin (p=0.040), and αSyn/TH (p=0.026) between 1.5 and 6 months.

**Figure 3B.** A main effect of age on the number of aggregates (clusters) was identified (F(2,6)=7.567, p=0.023) by one-way ANOVA, and multiple comparisons with Bonferroni correction revealed statistically significant difference between 1.5 and 12 months of age (p=0.03). No differences in the aggregate median size or number of localizations per cluster were found.

**Figure 3C.** Analysis of cluster size distribution has shown statistically significant increase in the number of medium-size aggregates - one way ANOVA revealed a main effect of age on the abundance of 100-300 nm (F(2,6)=9.784, p=0.013) and 300-500 nm (F(2,6)=7.668, p=0.022) clusters, and an increase between 1.5 and 12 months of age in these populations of clusters was revealed by multiple comparisons with Bonferroni correction (p=0.014, p=0.024 for 100-300 nm and 300-500 nm clusters, respectively). Direct comparison with t-test revealed also a statistically significant difference between 1.5 and 12 month-old animals for 20-100 nm clusters (p=0.013).

**Figure 4A.** Two-way ANOVA identified a main effect of age (F(2,31)=6.566, p=0.004) and a statistically significant interaction between genotype and age (F(2,31)=3.849, p=0.032). Multiple comparisons with Bonferroni correction revealed significant differences between C57Bl/6S and MI2 animals at 12 months of age (p=0.03) and between 3 and 12 month-old MI2 mice (p<0.001) and 6 and 12 month-old MI2 mice (p=0.019).

**Figure 4B.** At 3 months of age no difference between MI2 and C57Bl/6S mice was observed. At 6 months, interaction between genotype and sample time was identified by two-way mixed ANOVA (F(6,48)=3.470, p=0.006). In MI2 mice, following K^+^ stimulation, DA release was reduced compared to C57Bl/6S (t-test for individual sampling time points: p=0.038 at 60 min., p=0.009 at 100 min.). In older mice this deficit was more prominent (at 9 months - genotype x sample time interaction, F(6,54)=8.965, p<0.001; t-test, p=0.013 at 60 min, p=0.015 at 80 min, p=0.015 at 100 min; at 12 months - genotype x sample time interaction, F(6,30)=17.322, p<0.001; t-test, p=0.011 at 60 min, p=0.008 at 80 min, p=0.028 at 100 min).

**Figure 5B.** A main effect of genotype (F(1,24)=24.981, p<0.001) was identified by two-way ANOVA, and a significant reduction of TH+ neurons in MI2 mice compared to C57Bl/6S controls at 12 (p=0.018) and 20 (p<0.001) months of age, and between 9- and 20 months-old MI2 animals (p=0.015) were found using multiple comparisons with Bonferroni correction

**Figure 5E.** A main effect of genotype (F(1,18)=9.967, p<0.005) was identified by two-way ANOVA, and significant differences between C57Bl/6S and MI2 mice at 12 months of age (p=0.003), and between 9 and 12 month-old MI2 animals (p=0.017) were found using multiple comparisons with Bonferroni correction.

**Figure 6A.** A main effect of age on the rotarod performance was identified by two-way ANOVA (F(3,108)=7.299, p<0.001). Multiple comparisons with Bonferroni correction revealed statistically significant difference between 6 and 20 months (^###^p<0.001) and 12 and 20 months (^‡‡‡^p<0.001) in MI2 mice, and between MI2 and C57Bl/6S animals at 20 months (p=0.012).

**Figure 6B.** No statistically significant differences between MI2 and C57Bl/6S mice were identified using 25 mm rod (t-tests, p=0.082 and p=0.103, orientation time and transit time, respectively). No difference in orientation time between experimental groups was found using 15 mm rod (t-test, p=0.096), but there was a statistically significant difference between MI2 and C57Bl/6S animals in transit time on 15 mm rod (p=0.028, t-test).

**Figure 7D.** Statistically significant differences between anle138b- and placebo-treated MI2 mice were identified by t-test for inner cluster density (p=0.031) and percentage of non-clustered 1-120 hαSyn (p=0.005).

**Figure 7E.** Statistically significant differences between anle138b- and placebo-treated MI2 mice were identified by t-test for high molecular weight (p=0.005) and monomeric (p=0.009) 1-120hαSYN.

**Figure 8A.** Three-way mixed ANOVA identified an effect of genotype (F(1,17)=10.733, p<0.001), effect of sampling time (F(6,102)=41.756, p<0.001), two-way genotype x sampling time interaction (F(6,102)=8.416, p<0.001) and three-way interaction between sampling time, genotype and treatment (F(6,102)=2.355, p=0.036). Two-way ANOVA run for individual sampling time points identified a main effect of genotype at 60 min (F(1,18)=10.468, p=0.005), 80 min (F(1,18)=12.126, p=0.003) and 100 min (F(1,18)=10.146, p=0.005), and two-way genotype x treatment interaction at 80 min (F(1,18)=7.537, p=0.013). Multiple comparisons with Bonferroni correction revealed statistically significant differences between placebo-treated MI2 and C57Bl/6S mice at 60, 80 and 100 min (p=0.004, p<0.001, p=0.003, respectively) and between placebo- and anle138b-treated MI2 mice at 60, 80 and 100 min (p=0.048, p=0.018, p=0.038, respectively).

**Figure 8C.** A main effect of genotype was identified by two-way ANOVA (F(1,8)=11.682, p=0.009). Multiple comparisons with Bonferroni correction revealed statistically significant difference in number of nigral DA neurons between placebo-treated C57Bl/6S and MI2 mice (p=0.005), but not between anle138b-treated animals (p=0.365). There was also a significant difference between placebo- and anle138b-treated MI2 mice (p=0.023).
